# Ataxia Telangiectasia patient-derived neuronal and brain organoid models reveal mitochondrial dysfunction and oxidative stress

**DOI:** 10.1101/2024.01.29.577683

**Authors:** Hannah C Leeson, Julio Aguado, Cecilia Gómez-Inclán, Harman Kaur Chaggar, Atefah Taherian Fard, Zoe Hunter, Martin F Lavin, Alan Mackay-Sim, Ernst J Wolvetang

## Abstract

Ataxia Telangiectasia (AT) is a rare genetic disorder caused by mutations in the *ATM* gene and is characterized by oxidative stress, premature ageing, and progressive neurodegeneration of the cerebellum. The molecular mechanisms driving the neurological defects AT remain unclear, mainly due to lack of human neuronal models. Here, we use AT patient-derived pluripotent stem cells (iPSCs) and iPSC-derived neurons and brain organoids to comprehensively explore mitochondrial dysfunction, oxidative stress, and senescence phenotypes. We identified mislocalisation of mitochondria, a prevailing reduction in mitochondrial membrane potential, and increased oxidative stress in AT patient-derived iPSC and neuronal cultures that was restored by ATM gene correction. Cortical brain organoids from AT patients also display extensive oxidative stress, increased levels of senescence, and impaired neuronal function that could be counteracted with antioxidant treatment. Transcriptomic analysis identified disruptions in regulatory networks related to mitochondrial function and maintenance, including alterations in the PARP/SIRT signalling axis and dysregulation of key mitophagy and mitochondrial fission-fusion processes. Our study reveals that progressive mitochondrial dysfunction and aberrant ROS production are hallmarks of AT, and lead us to conclude that ATM is a master regulator of mitochondrial homeostasis.

## Introduction

Ataxia Telangiectasia (AT) is a rare autosomal recessive disorder caused by mutations in the ataxia-telangiectasia-mutated (*ATM*) gene, which encodes a 350 kDa serine/threonine kinase involved in DNA damage response and antioxidant defence pathways. Symptoms include immune defects, increased risk of cancers, premature aging, and ataxia resulting from progressive neurodegeneration of the cerebellum [1, 2]. In a healthy individual, ATM is activated in response to double strand DNA breaks and functions as a master regulator of a complex signalling axis that coordinates DNA repair, cell cycle arrest or apoptosis [3, 4]. Although neurons do not undergo DNA replication, double strand DNA breaks can still arise from both physiological neuronal activity [5], and from a mitochondria-to-nucleus retrograde signalling cascade that originates with dysfunctional mitochondria and generation of reactive oxygen species [ROS; 6]. Nevertheless, it remains unclear how defective ATM leads to degeneration of post-mitotic cells in the cerebellum and hippocampus of AT patients, and debate continues as to whether this is simply attributable to a faulty DNA damage response [7], or if other processes are at play, such as improper cell cycle re-entry [8], oxidative damage [9] or impaired synaptic functions [10].

ATM emerged as an important modulator of mitochondrial homeostasis and oxidative stress following the identification of increased levels of ROS in the cerebellum of ATM^-/-^ mice [11–13]. Elevated oxidative stress and perturbed mitochondrial function has also been detected in human AT patients [14], in lymphocytes and immortalised lymphoblastoid cells [15], as well as in AT patient fibroblasts, which displayed decreased mitophagy [16]. This is of particular interest when considering the important role mitochondria play in neuronal function, and the particular sensitivity of the cerebellum to mitochondrial impairment [17, 18], where any deviation to mitochondrial homeostasis can negatively impact neuronal integrity and function [19]. ATM localises to the mitochondria and may be directly activated by oxidative stress in a mitochondrial dependent manner and in the absence of DNA damage response pathway activation, demonstrating that ATM also functions as a sensor of ROS and mediates antioxidant responses [16, 20, 21]. Oxidized ATM forms a disulfide–cross-linked dimer [20], and when activated by oxidative stress, ATM has over 2500 protein targets [22], compared with 700 protein targets identified in response to DNA damage [23].

Despite this, there has been a limited number of studies dedicated to examining mitochondrial dysfunction in non-transformed human neuronal models. Although ATM-deficient mice recapitulate some of the cellular defects observed in AT, other AT-related defects such as neuronal degeneration are not evident in ATM^-/-^ mice [24]. Neuronal models derived from induced pluripotent stem cells (iPSCs) of individuals with AT therefore present an opportunity to examine the role of ATM in mitochondrial dysfunction and oxidative stress in a human system. Here we examine mitochondrial content, membrane potential and oxidative stress levels in AT patient neural models across different stages of maturation, from primary tissues, through iPSC and iPSC-derived neuronal progenitors, 2D neuronal cultures and finally brain organoids. We observe significant levels of oxidative stress that compound with neuronal maturation and highlight several mechanisms that putatively contribute to neuronal degeneration in AT.

## Results

### Primary olfactory epithelial cells from AT patients do not exhibit mitochondrial disturbances

To investigate disease specific differences in mitochondrial function in primary tissue from AT and control patients, we utilised human olfactory epithelial (ONS) cells. ONS cells are obtained from the primary olfactory epithelial mucosa, where olfactory receptor neurons are regenerated from a resident progenitor pool [25]. They possess some characteristics of neural epithelial cells [26] and exhibit disease-specific differences in several neurological disorders including AT [26, 27]. ONS cells (Sup Fig 1A) from five AT patients were confirmed to have little to no expression of ATM protein (Sup Fig 1B) compared to healthy age matched controls, and were characterised by immunochemistry as Nestin+, PAX6–, βIII tubulin–, and Map2– (Sup Fig 1C). We quantified total mitochondrial content with MitoTracker Deep Red (Sup Fig 1D) and mitochondrial membrane potential by TMRE labelling (Sup Fig 1E), which is indicative of energy production and respiratory chain function respectively [28]. Reactive oxygen species (ROS) production was measured using MitoTracker Red CM-H_2_Xros; a reduced, non-fluorescent MitoTracker that fluoresces upon oxidation by ROS species prior to mitochondrial sequestration (Sup Fig 1F). Our unbiased analysis pipeline (Sup Fig 1G) revealed no significant differences in any of these parameters between control and AT ONS cells. Mitochondrial content and function were further interrogated by western blot for individual oxidative phosphorylation (OXPHOS) complexes, and no significant differences were observed between control and AT (Sup Fig 2). We concluded that ONS cells from AT patients do not exhibit altered mitochondrial content, mitochondrial respiratory complex expression, membrane potential or mitochondrial ROS production as compared to control ONS cells.

### Mitochondrial perturbances emerge in AT stem and neuronal progenitor cell populations

Given that neuronal degeneration is one of the most debilitating aspects of AT, and currently one of the least understood, we extended our studies to investigate mitochondrial dysfunction in a relevant neuronal model. Hence, we reprogrammed ONS cells to iPSCs with the aim of investigating mitochondrial dysfunction in patient-derived neurons and brain organoids. AT ONS line 3 and control ONS were reprogrammed into iPSCs as described previously [29, 30]. These were used in subsequent experiments in addition to our previously generated AT32 patient and isogenic (gene corrected) iPSC pair [31–33]. Both AT iPSC lines have mutations resulting in a truncated non-functional ATM protein [Sup Fig 3; 29, 32]. Western blot confirmed reduced ATM expression in AT ONS derived-iPSCs and AT32 mutant iPSC clonal lines, in agreement with their respective ONS expression levels. AT32 gene correction rescued expression to levels comparable with control ONS derived-iPSCs (Fig 1A).

**Figure 1:**
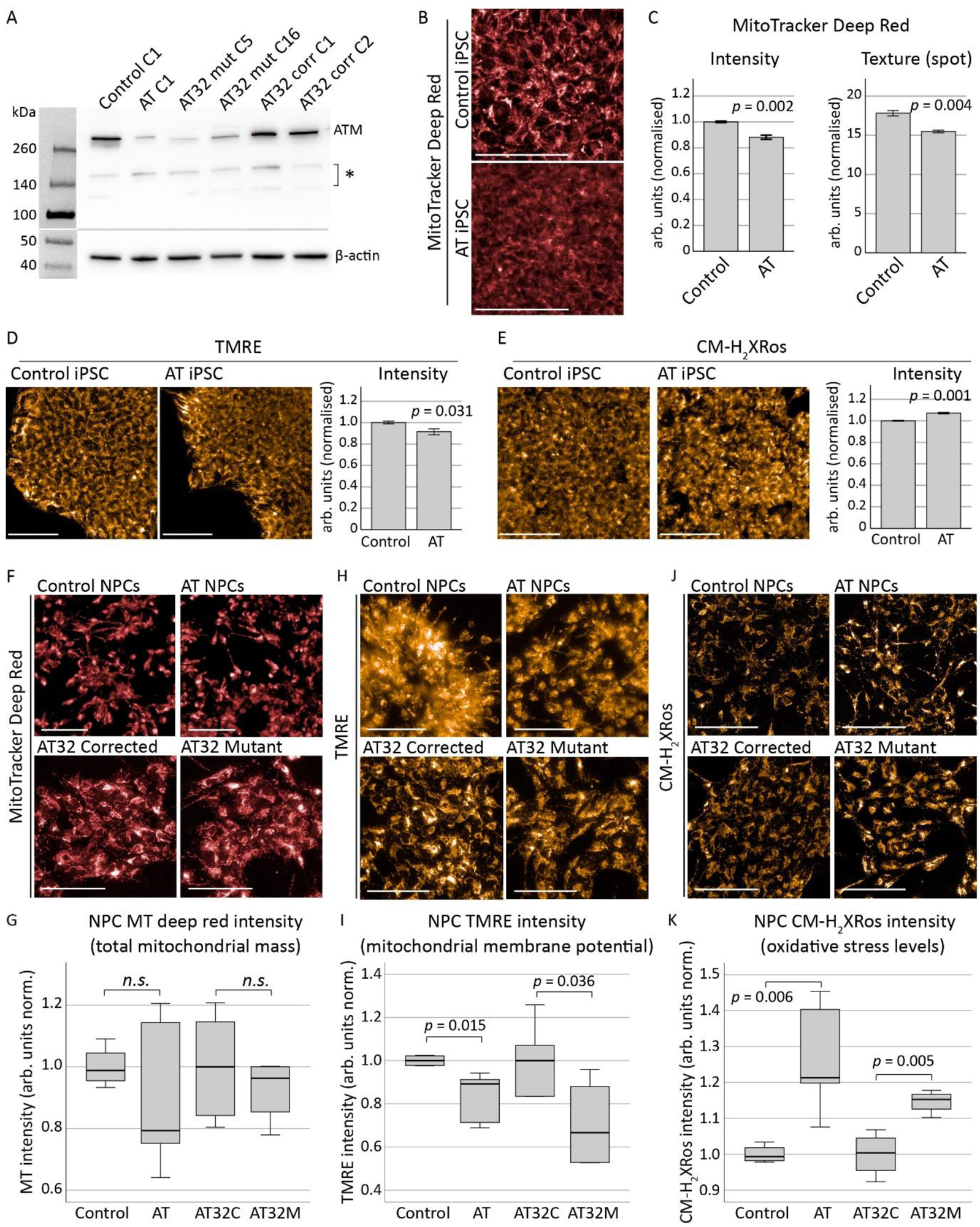
Mitochondrial assessments on AT patient-derived iPSCs and NPCs. **A)** Western blot confirmed a reduction in ATM protein levels in the AT ONS-derived iPSC line (AT) compared to the control line, as well as in two clones of AT32 iPSC lines. Gene correction in the AT32 line restored ATM expression to levels similar those seen in the control lines. Asterisk indicates non-specific bands. C refers to the clone number. **B-E)** Mitochondrial content and function in AT and control iPSC lines were assessed by MitoTracker Deep Red and TMRE, while oxidative stress was measured by CM-H_2_Xros. MitoTracker Deep Red **(B,C)** identified significant decreases in mitochondrial content (t(4) = -7.026, *p* = 0.002) and texture analysis of staining demonstrated AT iPSCs contained structurally altered mitochondria compared to the mitochondria of the control iPSC line (t(4) = -6.106, *p* = 0.004), using an unbiased algorithm to quantify saddle/spot/ridge variables of each puncta. TMRE **(D)** demonstrated a significant reduction in membrane potential (t(8) = -2.854, *p* = 0.031) in AT iPSCs compared to control iPSCs, and an increase in oxidative stress levels as measured by CM-H_2_Xros **(E)** was also observed in AT iPSCs (t(4) = 8.947, *p* = 0.001). Error bars are mean ± 1 SE. **F-K)** AT, control and AT32 iPSCs were differentiated to NPCs and assessed for mitochondrial content, function and oxidative stress. MitoTracker Deep Red **(F-G)** identified no significant difference in mitochondrial content between control and AT NPCs (t(8) = 1.124, *p* = 0.303), nor between AT32 mutant and gene corrected NPCs (t(8) = 0.797, *p* = 0.449). TMRE **(H-I)** demonstrated a significant reduction in membrane potential in both AT and AT32 mutant lines compared to the control and AT32 corrected NPCs (t(8) = -3.411, *p* = 0.015 and t(8) = 2.512, *p* = 0.036 respectively). **J-K)** Oxidative stress levels as measured by CM-H_2_Xros were significantly increased in both AT lines (AT v control; t(8) = -4.384, *p* = 0.006, AT32 pair; t(6) = -4.263, *p* = 0.005]. Box; median ± IQ, and whiskers; min and max data points. All scale bars indicate 100 µm. AT; AT ONS-derived iPSC line. Control; control ONS-derived iPSC line. AT32C; AT32 gene corrected iPSC line. AT32M; AT32 mutant iPSC line.

Mitochondrial content, membrane potential and oxidative stress in ONS-derived AT and control iPSC lines were next quantified (Sup Fig 3D). Significant decreases in both MitoTracker and TMRE were identified in AT iPSCs compared to control iPSCs (Fig 1B-D). Texture analysis of MitoTracker Deep Red staining revealed AT mitochondria exhibit altered structure and/or sub-cellular networking compared to the mitochondria of the control iPSC line. A significant increase in oxidative stress levels was also observed in AT iPSCs (Fig 1E), in agreement with previous observations [32]. ONS-derived AT and control iPSCs and the AT32 gene corrected (isogenic) iPSC pair were next differentiated to neural progenitor cells (NPCs) and neurons [34], and were confirmed to express comparable levels of progenitor and early neuron markers (Sup Fig 4). Mitochondrial content, membrane potential and oxidative stress were assessed as above. Image quantification revealed no significant difference in mitochondrial content between control and AT NPCs (Fig 1F, G), whereas membrane potential in both ONS-derived AT NPCs and AT32 mutant NPCs was significantly reduced compared to their respective controls (Fig 1H, I). Furthermore, oxidative stress levels were increased in ATM deficient NPC cultures with the mean CM-H_2_Xros fluorescence intensity increasing by 26% in ONS-derived AT and 15% in AT32 mutant NPCs compared to their control counterparts (Fig 1J, K).

We concluded that AT iPSC have less and structurally altered mitochondria and exhibit a modest increase in ROS production as compared to control counterparts. Further, AT iPSC do not show defective differentiation into NPCs and that unlike iPSC cultures, NPCs do not have abnormal mitochondrial content, but do exhibit a reduction in mitochondrial membrane potential and increased ROS production as compared to unrelated and isogenic controls in agreement with undifferentiated iPSCs. Given these significant defects in mitochondrial membrane potential and increased ROS production, we hypothesized that this would likely have downstream impacts on mitochondrial homeostasis during neuronal maturation.

### AT patient iPSC-derived neurons show defective and mis-localised mitochondria

Both AT neurons and their respective control/isogenic corrected neurons were matured for 2-4 weeks (Fig 2A), and by 2 weeks >90% of cells expressed βIII tubulin and Map2 neuronal markers (Fig 2B). AT mutant neurons displayed evidence of blebbing and fragmentation consistent with neuronal degeneration, which was most apparent following βIII tubulin staining. We assessed mitochondrial content and function in both 2- and 4-week-old neurons to monitor changes in mitochondrial status over the course of maturation. Fluorescence intensities of mitochondrial dyes were measured in both the soma (neuronal cell body) as well as in the neuronal processes (neurites; Sup Fig 5A). 2-week-old AT neurons exhibit perturbances in mitochondrial content and localisation, with reduced cytoplasmic MitoTracker intensity and an increase in intensity in the neurites (Sup Fig 5B, C), suggesting mitochondria are becoming localised to the neuron periphery. Mitochondrial membrane potential correlated with MitoTracker observations (Sup Fig 5D, E). In contrast to NPC cultures, 2-week neurons showed no significant differences in oxidative stress levels between AT32 mutant and isogenic corrected neurons (Sup Fig 5F, G).

**Figure 2:**
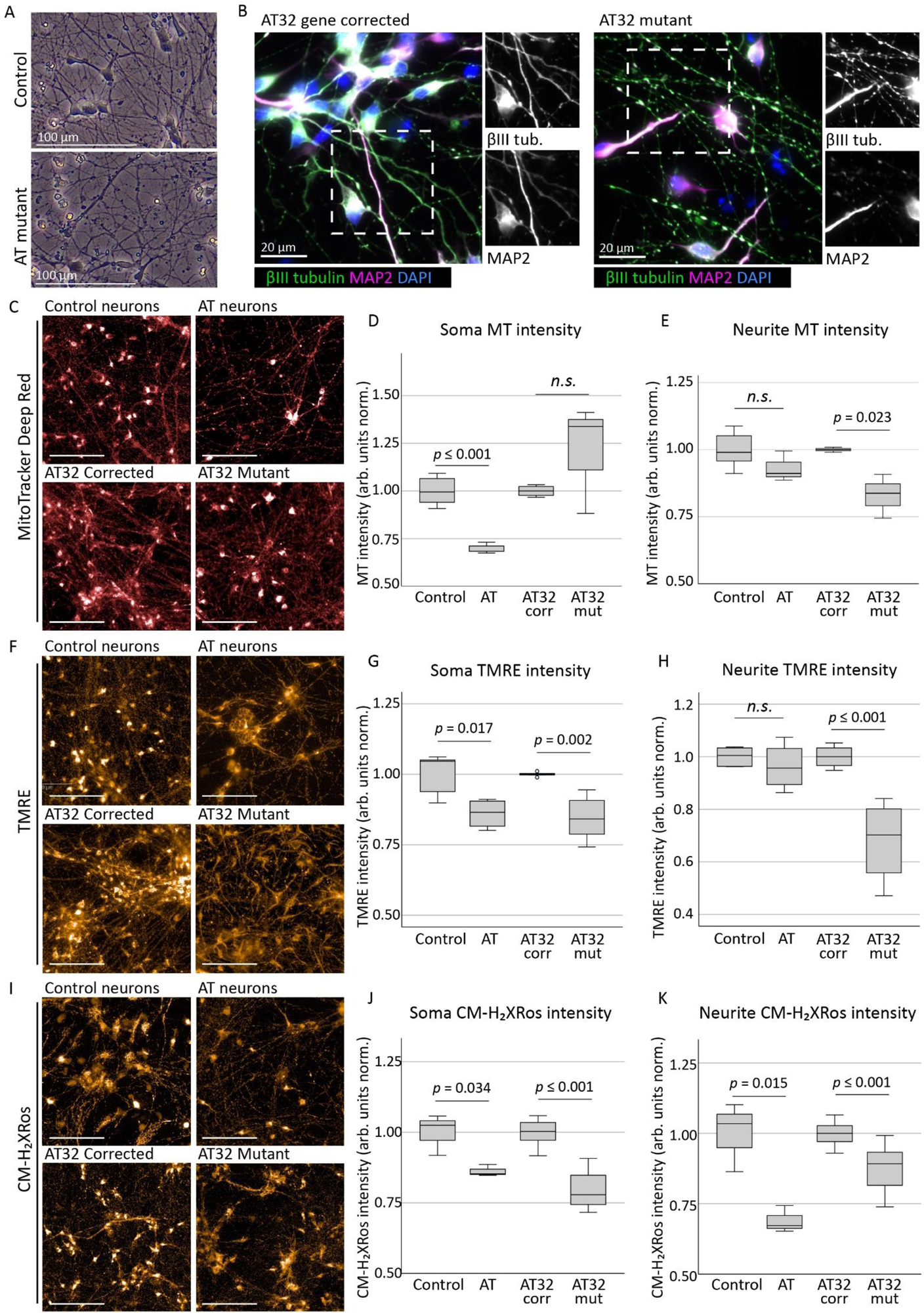
Mitochondrial assessments on AT patient iPSC-derived neurons. **A)** Neuronal networks generated from AT patient and control/gene corrected iPSCs were cultured on PLO and laminin and matured for 2-4 weeks. **B)** Neurons expressed neuronal markers βIII tubulin and MAP2, with AT mutant neurons displaying degrees of fragmentation consistent with a neurodegenerative phenotype. **C)** Mitochondrial content in AT patient and control/gene corrected neurons were assessed by MitoTracker Deep Red. Soma intensity **(D)** showed a significant reduction in AT neurons compared to control (t(6) = -6.177, *p* ≤ 0.001), though not in AT32 neurons (t(6) = -1.269, *p* = 0.165). Conversely, neurite fluorescence levels **(E)** identified a significant reduction in mitochondria localised to the projections in AT32 mutant neurons (t(4) = 3.566, *p* = 0.023) while the AT and unrelated control lines were not altered (t(9) = 1.645, *p* = 0.134). TMRE assessment **(F)** demonstrated consistent impairments in mitochondrial membrane potential in ATM deficient neurons at 4 weeks, particularly in soma mitochondria (**G;** AT v control t(7) = -3.100, *p* = 0.017; and AT32 isogenic lines t(11) = 5.285, *p* = 0.002). Neurite fluorescence of TMRE **(H)** was also reduced in AT32 mutant neurons (t(11) = 6.102, *p* ≤ 0.000), though not in the AT neuron line vs its unrelated control (t(7) = 0.853, *p* = 0.422). CM-H_2_Xros staining **(I)** showed reduced soma fluorescence intensity **(J)** in AT (t(4) = 3.163, *p* = 0.034) and in AT32 (t(35) = 10.987, *p* ≤ 0.000) neurons compared to their respective controls, as well as in neurite intensity **(K;** t(4) = 4.085, *p* = 0.015, and t(35) = 5.822, *p* ≤ 0.000, respectively). Scale bars 100 µm. Boxplots show the median ± IQ, with minimum and maximum values indicated by the whiskers.

By 4 weeks of maturation, neuronal cultures demonstrated differing mitochondrial phenotypes to their younger 2-week-old counterparts. ONS-iPSC derived AT neurons demonstrated impairments in mitochondrial localisation and/or trafficking, as indicated by decreased total mitochondrial staining in the cytoplasmic region (Fig 2C, D) and a 50% increase in the number of mitochondrial puncta in their neurites (Sup Fig 7A), while neurite spot intensities remained unchanged. We speculate that this may be related to impairments in mitochondrial turnover and recycling of damaged mitochondria, correlating with ATM’s role as a regulator of mitophagy [16, 35, 36] and of mitochondrial fission [37]. Conversely, AT32 mutant neurons had significant reductions in neurite mitochondria while retaining a consistent puncta count (Fig 2C, E, Sup Fig 6A). This also points to impairments in structural arrangements and fission/fusion of mitochondria, though it is likely that both scenarios are present and contribute to an overarching phenotype of mitochondrial impairment, with the discrepancies between cells lines likely reflective of patient-specific differences. TMRE continued to reveal perturbances in mitochondrial membrane potential in 4-week neuronal cultures (Fig 2F), in keeping with both NPCs and 2-week neurons. This was particularly evident in the cytoplasmic mitochondrial fluorescence levels (Fig 2G). Mitochondria localised to the neuronal processes in the AT32 mutant neurons also had significantly reduced membrane potential when compared to their isogenic AT32 corrected control neurons (Fig 2H).

Interestingly, when we assessed oxidative stress levels using MitoTracker CM-H_2_Xros in 4-week-old neuronal cultures we observed a significant decrease in fluorescence levels, contrary to observations in iPSC and NPC cultures (Fig 2I-K). Though unexpected, this reversal in fluorescence levels from that seen in stem and progenitor cell cultures is consistent with 2-week neuron observations, where no significant difference was observed. Hydrogen peroxide treatment in a control neuron culture was used as a positive control for oxidative stress (Sup Fig 6B), and we confirmed oxidisation and sequestration of CM-H_2_Xros dye into mitochondria occurred in a linear manner in both ATM deficient and gene corrected lines (Sup Fig 6C). Protein expression levels of individual OXPHOS complexes were quantified to determine if there were deficits that may contribute to impaired membrane potential. Complex III protein levels were significantly increased in AT32 mutant neurons, and other complexes were altered but did not reach significance (Sup Fig 6D-F).

While MitoTracker CM-H_2_Xros detects cellular ROS levels by fluorescing only once the dye becomes oxidised, mitochondrial sequestration is dependent upon membrane potential. We concluded that the loss of ATM in neurons results in chronic impairment of mitochondrial membrane potential, as indicated by TMRE, preventing CM-H_2_Xros uptake in mutant neurons. To further explore the oxidative stress phenotype, we cultured control neurons with ATM kinase inhibitors KU-55933 and KU-60019 prior to mitochondrial assessment at 4 weeks and interrogated mitochondrial content (MitoTracker Deep Red; Fig 3A-C), membrane potential (TMRE; Fig 3D-F) and oxidative stress (CM-H_2_Xros; Fig 3G-I). KU-60019 treatment resulted in a significant increase in cytoplasmic intensity of MitoTracker Deep Red, and mitochondrial puncta localised to the neurites also fluoresced brighter and were aggregated in appearance, indicative of mitochondrial fusion or decreased mitophagy as was previously reported [16]. Supporting this, a significant reduction in mitochondrial puncta localised to the neurites was observed following KU-55933 and KU-60019 exposure (Fig 3J). Increased TMRE fluorescence in both cytoplasmic and neurite mitochondria was observed following acute ATM inhibition (Fig 3D-F). This potentially indicates a higher rate of oxidative phosphorylation and may indicate a homeostatic or compensatory response downstream of acute ATM inhibition. In contrast to observations that ATM-deficient neurons showed decreased CM-H_2_Xros fluorescence due to chronically impaired mitochondrial membrane potential, we found that acute chemical inhibition of ATM in control neurons with otherwise healthy mitochondria resulted in a significant increase in CM-H_2_Xros fluorescence in mitochondria localised to both the cytoplasm and neurites (Fig 3G-I). This indicates there are significant levels of oxidative stress in neurons lacking a functional ATM signalling pathway and supports ATM’s role as a sensor of oxidative stress as well as in orchestrating the cellular response to ROS [20]. Notably, treatment with ATM inhibitors did not reduce the number of neurons based on nuclei count (Fig 3K), suggesting that ATM signalling deficiency drives mitochondrial defects in the absence of cytotoxic effects.

**Figure 3:**
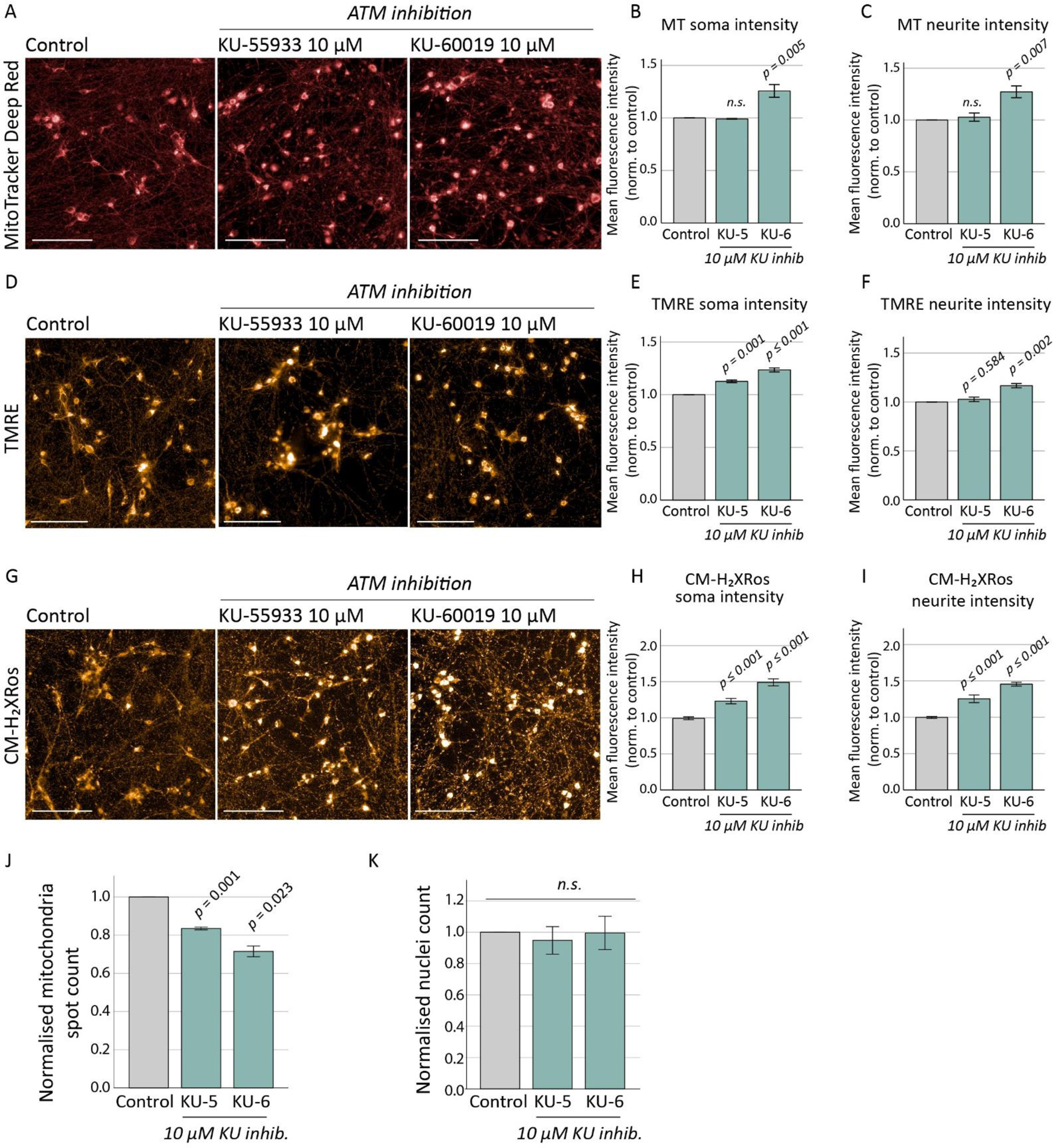
Mitochondrial assessments on control neurons treated with chemical ATM inhibitors. Control neurons were treated with ATM inhibitors KU-55933 and KU-60019 (10 µM) for 10 days prior to mitochondrial assessment at 4 weeks. Neurons were stained with MitoTracker Deep Red **(A)**, and KU-60019 treatment caused increases in fluorescence intensity in both soma [**B;** F(2, 8) = 19.001, *p* = 0.003] and neurite puncta [**C;** F(2, 8) = 13.918, *p* = 0.006] as determined by ANOVA, with Tukey post hoc analysis identifying *p* values of 0.005 and 0.007 respectively. KU-55933 made no significant differences to mitochondrial content. **D)** TMRE staining to measure membrane potential identified an increase in fluorescence levels in both soma and neurite staining following KU-55933 and KU-60019 treatments. **E)** Soma TMRE intensity: F(2, 8) = 78.289, *p* ≤ 0.001; Tukey post hoc analysis KU-55933 *p* = 0.001, and KU-60019 *p* ≤ 0.001 compared to control. **F)** Neurite TMRE intensity: F(2, 8) = 23.535, *p* = 0.001, Tukey post hoc analysis KU-55933 *p* = 0.584 and KU-60019 *p* = 0.002. **G)** CM-H_2_XRos fluorescence increased following chemical inhibition of ATM signalling, indicating a significant oxidative stress phenotype. **(H)** Soma CM-H_2_XRos fluorescence intensity: F(2, 18) = 49.530, *p* ≤ 0.001; Tukey post hoc analysis: *p* ≤ 0.001 and *p* ≤ 0.001 for KU-55933 and KU-60019 respectively, and **(I)** neurite CM-H_2_XRos fluorescence intensity: F(2, 18) = 33.655, *p* ≤ 0.001; Tukey post hoc analysis: *p* ≤ 0.001 and *p* ≤ 0.001 for KU-55933 and KU-60019 respectively. **J)** ATM chemical inhibition resulted in a significant reduction in number of mitochondrial puncta located in neuronal processes [F(3, 11) = 28.117, *p* ≤ 0.001] for both KU-55933 (*p* = 0.001) and KU-60019 (*p* = 0.023), though ATM inhibition did not affect the number of neurons based on nuclei count [**K;** F(2, 8) = 0.617, *p* = 0.571]. Scale bars; 100 µm. Error bars are mean ± 1 SE. KU-5; KU-55933. KU-6; KU-60019.

Collectively, we conclude that ATM-deficient neurons display severe mitochondrial dysfunction that become more pronounced over the course of the maturation process. The consistent reduction in mitochondrial membrane potential (observed from iPSCs through NPCs to both 2- and 4-week neuronal stages) is indicative of impaired electron transport chain dysfunction and may impact ATP synthase activity. Comparison of CM-H_2_Xros fluorescence in AT patient-derived neurons to control neurons treated with ATM inhibitors provided important insight into the extent of mitochondrial dysfunction in ATM mutant neuronal populations, as acute ATM inhibition did not reduce membrane potential and increased CM-H_2_Xros accumulation in mitochondria, indicative of significant ROS production. This also highlights the limitations of single-method mitochondrial assessments and the importance of careful selection of disease models.

### Transcriptional investigations into AT patient-derived neurons and BOs reveal strong oxidative stress and mitochondrial phenotypes

To gain further insight into the possible mechanisms and consequences of this mitochondrial dysfunction, we performed RNA sequencing of AT32 mutant and isogenic corrected iPSC, 2D neuronal cultures (2-week-old) and 3D brain organoids (100-day-old). We confirmed similar neural differentiation capacity between mutant and corrected neurons (Sup Fig 7), and comparison of AT32 corrected and mutant neurons identified 1,836 DEGs (Fig 4A); comparatively, brain organoids had nearly 9,000 DEG (Fig 4B). This was explained by principal component analysis, where gene set space determined that ATM-deficient neurons and organoids were separable from transcriptomes of ATM-proficient counterparts (Sup Fig 8A, 9A). Approximately 63% of neuronal DEGs were also identified in the brain organoid data set (Sup Fig 8C). KEGG pathway analysis identified strong mitochondrial impairments in AT32 mutant brain organoids, including upregulation of oxidative phosphorylation and TCA cycle pathways, as well as increases in ROS, neurodegeneration, and cellular senescence. Conversely, synapse, axon guidance, and cell cycle signalling pathways were downregulated (Fig 4C). GO terms showed significant enrichment of oxidation-reduction, cellular respiration, and electron transport chain processes, as well as response to stress processes including ROS, oxidative and ER stresses, and increased cell and neuron death and apoptotic processes (Fig 4D). Consistent with a non-functional ATM protein, DNA and double-strand break repair were downregulated, as well as processes relating to neurogenesis, nervous system development, and synaptic transmission. Notably, mitochondrial and organelle organisation, fission, localisation and transport processes were also impacted, aligning with our observations of alterations in mitochondrial localisation between neuronal cell body and processes (Fig 4E). The top 10-15 GO and KEGG terms for all categories are shown in Sup Figs 8 and 9.

**Figure 4:**
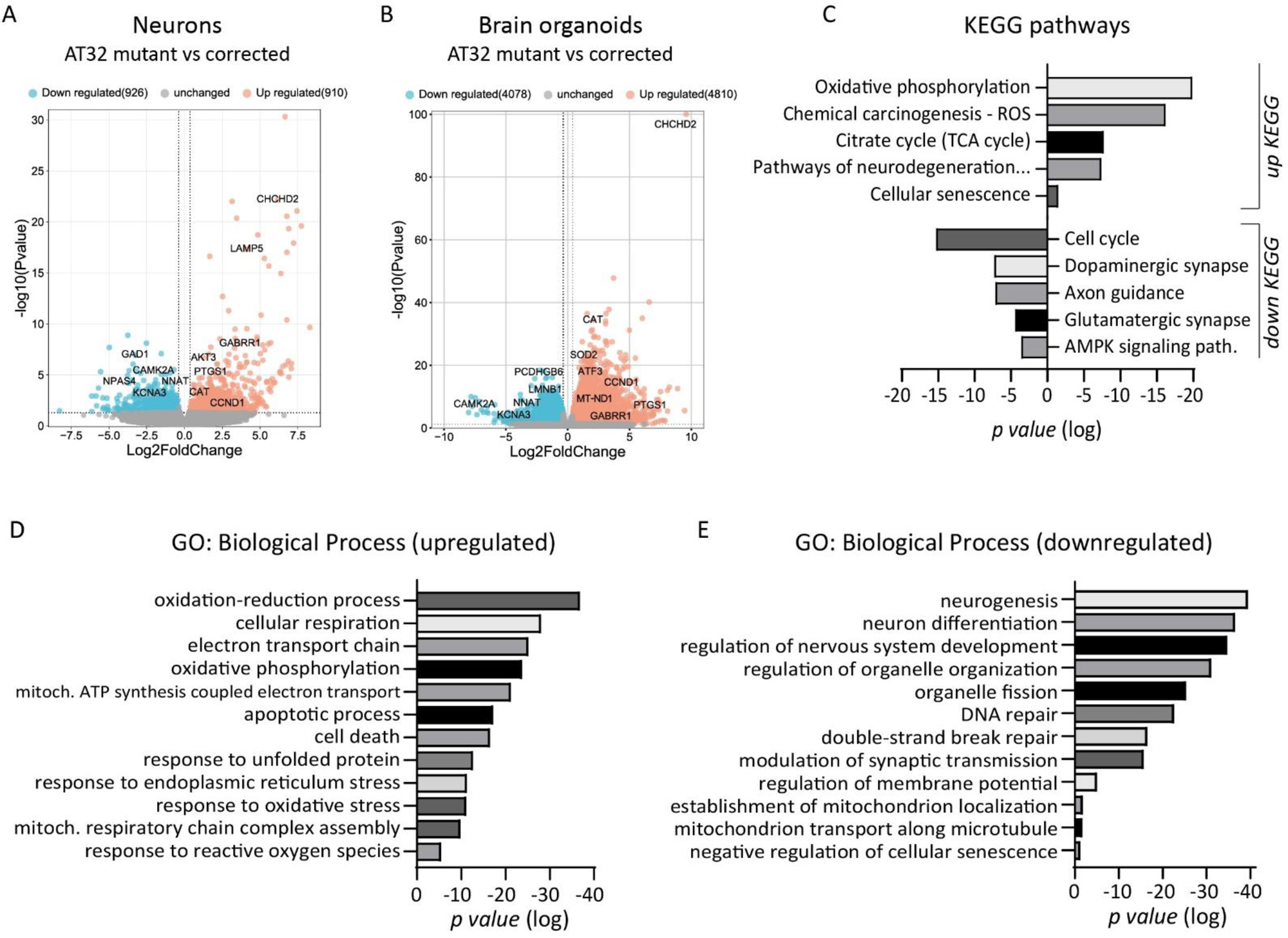
Transcriptomic analysis of AT32 neuron and brain organoid cultures. RNA sequencing was conducted on AT32 2-week-old neurons and 100-day-old brain organoids. **A)** Volcano plots showing comparison between AT32 mutant and gene corrected neurons identified 1836 DEGs (910 genes upregulated and 926 down regulated in the mutant compared to the corrected), while comparatively, 100-day-old brain organoids had nearly 9,000 DEG (**B;** 4810 genes upregulated and 4078 down regulated in the mutant compared to the corrected). KEGG pathway **(C)** and GO analysis **(D, E)** were conducted on up and down DEGs separately, and were enriched for pathways involved in mitochondrial processes, such as oxidative phosphorylation and electron transport chain, as well as response to stress processes including oxidative and ER stresses. DEG; differentially expressed gene. GO; gene ontology.

We next investigated the extent of oxidative stress in both neurons and organoids by analysing gene expression patterns of markers of oxidative stress (Fig 5A). Both AT32 mutant neurons and organoids demonstrated a distinct oxidative stress phenotype compared to the gene corrected, with a higher number of marker genes reaching significance in the organoid models, consistent with our earlier conclusion that mitochondrial dysfunction compounds with increasing neuronal maturity. One of the most notable genes upregulated in both ATM mutant neurons and organoids is *CHCHD2* (9.6 Log2FC, *p* = 3.3E-101; also known as *MNRR1*), a key regulator of oxidative and cell stress responses [38, 39], as well as mitochondrial metabolism and the electron transport chain [40], mitochondrial biogenesis and morphology [41, 42]. *CHCHD2* can also act as an inhibitor of apoptosis [43, 44], and is associated with neurodegeneration and Parkinsons disease [45–50]. Other notable oxidative stress genes upregulated in ATM mutant neurons and organoids include the superoxide dismutases *SOD2* and *SOD3, PTGS1* (cyclooxygenase 1; COX-1), *CAT* (CATALASE) and *SQSTM1* (p62). CATALASE and p62 have been linked to autophagy of peroxisomes mediated by ATM phosphorylation of PEX5, which recruits p62 and directs the autophagosome to peroxisomes to induce pexophagy in response to ROS [51]. ATM has previously been identified at the peroxisome [52], and contains a FATC domain thought to be a peroxisome targeting sequence. Interestingly, AT patients with mutations in this FATC domain do not display the radiosensitivity typical of AT patients [20] indicating that this domain functions independently of ATM’s DNA damage response.

**Figure 5:**
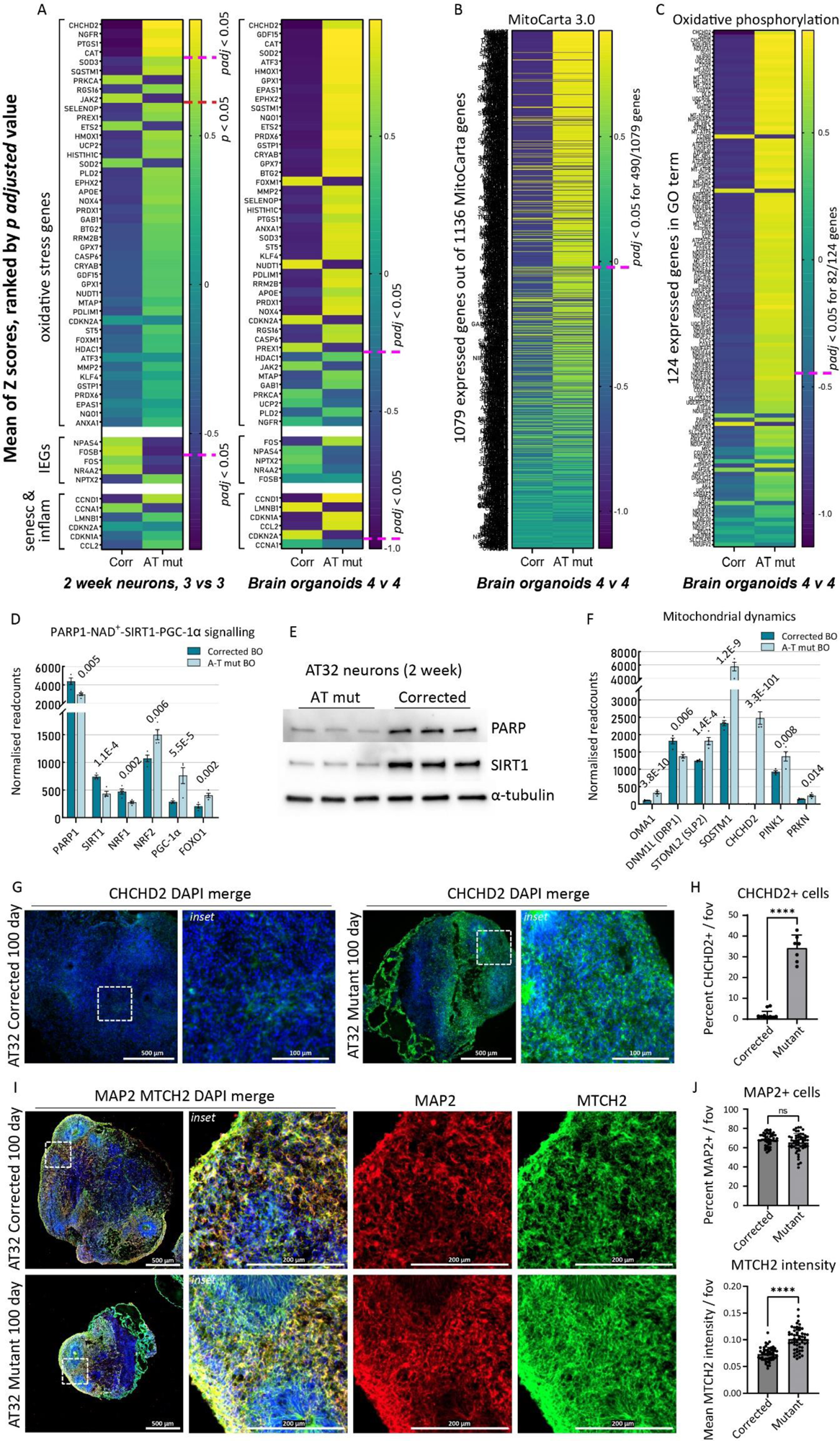
Oxidative stress signatures and mitochondrial perturbances revealed in AT32 neurons and brain organoids. **A-C)** The mean Z-scores were calculated from AT32 2-week-old neuron and 100-day-old brain organoid RNA sequencing data and were plotted as heatmaps for AT mutant and corrected, with genes ranked by their adjusted *p* value. The pink dashed line indicates where *padj* < 0.05 falls. **A)** A panel of oxidative stress markers, IEGs, and markers of senescence and inflammation were plotted comparing mutant and corrected neuron and brain organoids. **B)** The mean Z-scores of all expressed MitoCarta 3.0 genes were plotted comparing AT mutant and corrected brain organoids, as well as for **(C)** all expressed genes comprising the GO term ‘Oxidative Phosphorylation’. **D)** Normalised readcounts of genes in the PARP1-SIRT1 signalling pathway from brain organoid RNA sequencing data, including *PARP1*, *SIRT1*, *NRF1*, NRF2 (*NFE2L2*), PGC-1α (*PPARGC1A*) and *FOXO1*. *padj* values as indicated. **E)** Western blot analysis of PARP and SIRT1 expression in whole cell extracts from 3 independent AT32 2-week-old neuronal cultures. α-tubulin was used as loading control. **F)** Normalised readcounts of genes involved in fission/fusion and mitophagy from brain organoid RNA sequencing data, including *OMA1*, *DNM1L* (DRP1), *STOML2* (SLP2), *SQSTM1* (p62), *CHCHD2*, *PINK1* and *PRKN*. *padj* values as indicated. **G)** Representative immunofluorescence images of AT32 mutant and gene corrected brain organoid sections stained for CHCHD2. Scale bars 500 µm, inset 100 µm. **H)** AT32 mutant organoids had a significantly higher percentage of CHCHD2 positive cells [t(18) = 17.10, *p* ≤ 0.001], 7 to 10 FOV from 5 brain organoids. **I)** Representative immunofluorescence images of AT32 mutant and gene corrected brain organoid sections stained for MTCH2 and MAP2. Scale bars 500 µm, inset 200 µm. **J)** The percentage of MAP2 positive cells remained unchanged [t(105) = 1.978, *p* = 0.051] while MTCH2 intensity was increased in AT32 mutant organoids [t(105) = 8.466, *p* ≤ 0.001], 50 to 55 FOV from 5 brain organoids. Error bars mean ± SD. BO; brain organoid. FOV; field of view. Corr; AT32 gene corrected; Mut; AT32 mutant. IEG; immediate early gene. GO; gene ontology.

To indirectly gauge neuronal activity, we examined the expression of immediate early genes (IEGs), which are known to be dependent on functional mitochondria. We found that expression of *NPAS4*, *FOSB*, *FOS* and *NR4A2* were significantly reduced in ATM mutant neuronal cultures (Fig 5A), aligning with the downregulation of KEGG and GO terms relating to synaptic transmission and function (Fig 4). This was not evident in the organoid model system, presumably due to organoids containing numerous other cell types that can mask subtle changes in neuronal gene expression levels. Markers of senescence and inflammation were also assessed, since we have previously shown that AT brain organoids have increased senescence compared to embryonic stem cell derived controls [53]. In agreement with these data, a strong senescence phenotype was also observed in our 100-day old ATM mutant organoids compared to their isogenic controls.

By way of unbiased assessment, we analysed the mean gene expression for all genes included in the MitoCarta 3.0 catalogue (Fig 5B). This catalogue comprises a comprehensive list of 1136 human genes encoding proteins with mitochondrial localisation. We also analysed all expressed genes comprising the oxidative phosphorylation GO term (Fig 5C). The results showed significant mitochondrial impairment in ATM mutants, with enrichment of approximately half of the 1079 expressed MitoCarta3.0 genes, and two thirds of genes involved in oxidative phosphorylation, particularly ubiquinol-cytochrome C reductases (*UQCR* genes) and NADH-ubiquinone oxidoreductase subunits (*MT-ND*, *NDUF* genes). Consistent upregulation of these genes led us to believe they are compensatory mechanisms for a chronic reduction in NAD^+^ levels, and indeed AT null mice were previously found to exhibit depleted levels of NAD^+^ [54]. This NAD^+^ deficiency was downstream of poly(ADP-ribose) polymerase 1 (PARP1), and results in SIRT1 inactivation and subsequent defects in mitophagy and mitochondrial dysfunction. Hence, we investigated the PARP1-SIRT1 signalling pathway in our ATM mutant models, and included nuclear respiratory factor 1 (NRF1), a phosphorylation target of ATM [55] and NRF2 (regulated by ROS and/or ATM; [56, 57]), both of which are key transcription factors in mitochondrial biogenesis and antioxidant function [58], and which interact with PARP1, SIRT1, and transcriptional co-activators PGC-1α and FOXO1 [59, 60]. We found mRNA levels of *PARP1*, *SIRT1* and *NRF1* were decreased in the ATM mutant organoids, consistent with an NRF1 protein reduction in ATM null cells observed by [55] while *NRF2*, *PGC-1α* and *FOXO1* were increased (Fig 5D). In agreement with these observations, western blot analysis confirmed that protein levels of PARP1 and SIRT1 were significantly decreased in ATM mutant neurons (Fig 5E). We further hypothesised that the changes in upstream regulators of mitochondrial network dynamics (*CHCHD2*, *PARP1*, *PGC-1α*) would have downstream impacts on genes controlling fission/fusion and mitophagy. Indeed, we identified disturbances in key fission genes *OMA1*, *DNM1L* (DRP1) and *STOML2* (SLP2) [61] as well as significant upregulation in mitophagy genes *SQSTM1* (p62), *PINK* and *PRKN* (Fig 5F, Sup Fig 10A). Next, we confirmed that upregulated *CHCHD2* expression in ATM mutant organoids was reflected by protein expression levels, observing a significant increase in CHCHD2 immunofluorescence in AT32 mutant organoid sections compared to isogenic controls (Fig 5G, H). We also assessed whether the transcriptional upregulation of mitochondrial genes in mutant organoids correlated with altered mitochondrial content, by immunolabelling AT32 organoid sections with the mitochondrial outer membrane protein MTCH2 (Fig 5I). MTCH2 intensity was significantly increased in the mutant organoids, while the percentage of MAP2 positive cells remained unaltered (Fig 5J).

### AT patient-derived neuronal models display inflammation and senescence features associated with oxidative stress and mitochondrial dysfunction

A significant aspect of AT pathology and highly interconnected to mitochondrial dysfunction is cellular senescence; together they constitute two of the hallmarks of aging associated with AT [62]. Mitochondrial dysfunction is a prevailing feature and causative driver of cellular senescence [63], with aberrant fission and fusion, excessive ROS production, production of a pro-inflammatory secretory phenotype, and imbalances in mitochondrial metabolites, particularly NAD, shown to contribute to senescence phenotypes [64, 65]. We have previously identified premature senescence in AT ONS-iPSC derived brain organoids compared to unrelated controls [53], hence, we investigated the extent of senescence and inflammation in our AT32 mutant and isogenic organoids.

KEGG pathway analysis (Fig 4) indicated an increase in cellular senescence in ATM mutant brain organoids, and we analysed the mean expression for all transcribed genes comprising the Cellular Senescence GO term (Fig 6A). Nearly half of the genes comprising this GO term were significantly dysregulated. Normalised read count assessment of brain organoids identified classical markers of senescence, including increased levels of *CCND1*, *CDKN1A* (p21) and *CCL2* in mutant BOs, while *LMNB1* and *CDKN2A* (p16) were reduced (Fig 6B). 2-week-old neuronal cultures did not demonstrate significant differences (Fig 5A), and so to determine if senescence would become evident upon further neuronal maturation, we generated AT32 mutant and corrected neurons and matured them for 10 weeks. By contrast, we detected a 3-fold increase in the percentage of cells that exhibit senescence-associated β-galactosidase staining in AT mutant neurons (SA-β-Gal; Fig 6C, D). Significant increases in expression of p16 and p21 was observed via immunochemistry in the aged neuron cultures, confirming the increases in senescence levels in AT mutant neurons (Fig 6E-G).

**Figure 6:**
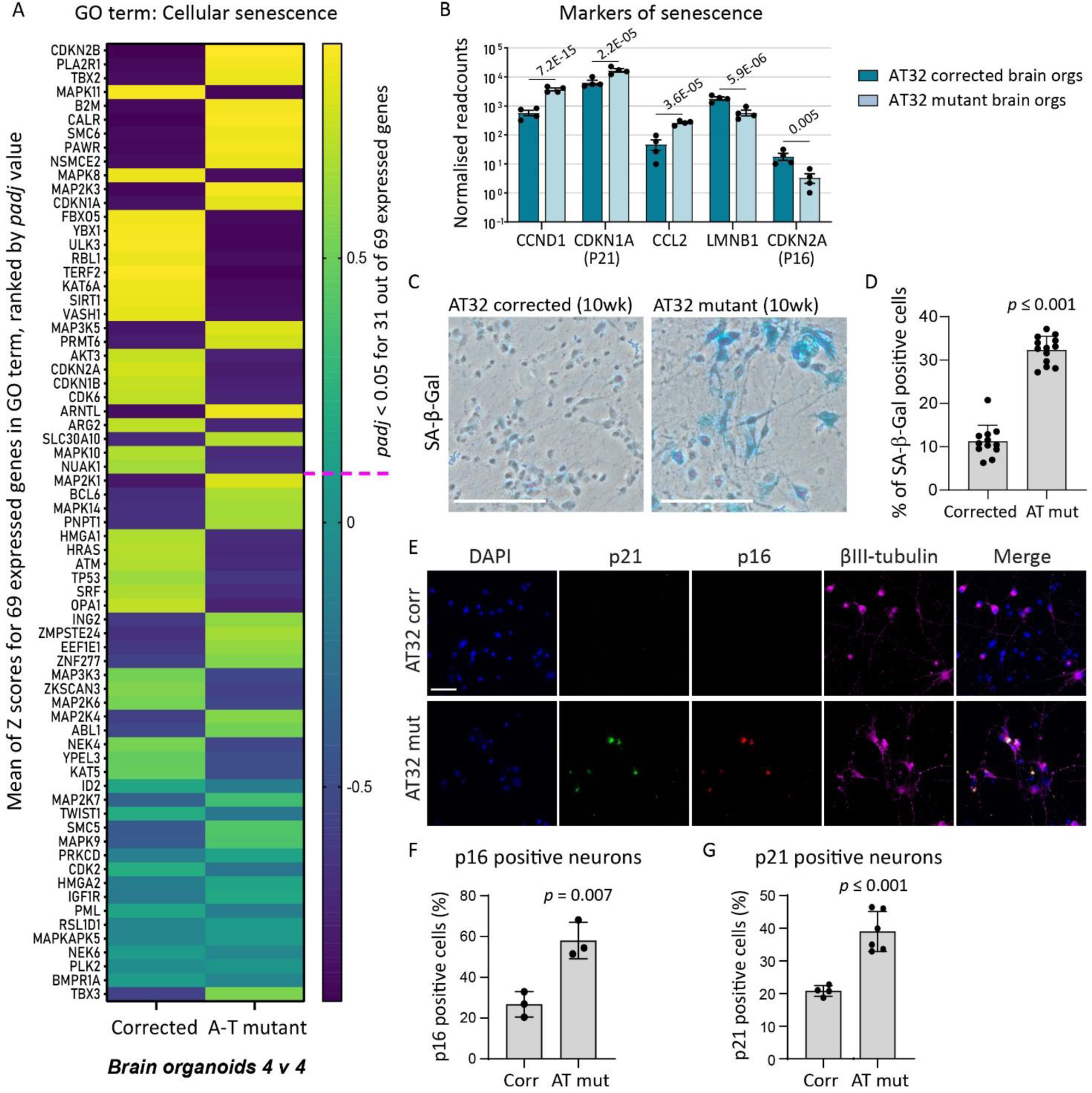
Senescence signatures revealed in AT32 neurons and brain organoids. **A)** The mean Z-scores of all expressed genes comprising the GO term ‘Cellular Senescence’ were plotted as a heatmap from AT32 brain organoid RNA sequencing data. Genes were ranked by their adjusted *p* value, and the pink dashed line indicates where *padj* < 0.05 falls. **B)** Normalised readcounts of markers of senescence from brain organoid RNA sequencing data, including *CCHD1*, *CDKN1A* (p21), *CCL2*, *LMNB1*, *CDKN2A* (p16). *padj* values as indicated. **C)** Representative images from SA-β-gal assays performed on AT32 mutant and corrected 10-week-old neurons. Scale bar 100 µm. **D)** AT32 mutant neurons had a significantly higher percentage of SA-β-gal positive staining, t(23) = 15.33, *p* ≤ 0.001. 12 FOV from 3 independent differentiations. **E)** Representative immunofluorescence images of AT32 mutant and gene corrected 10-week-old neurons stained for p21 and p16. Scale bar 100 µm. **F)** Quantification of p16 positive neurons [t(4) = 4.976, *p* = 0.007] and **G)** p21 positive neurons [t(8) = 5.680, *p* ≤ 0.001]. 3 independent differentiations. Error bars mean ± SD. FOV; field of view. Corr; AT32 gene corrected; Mut; AT32 mutant. GO; gene ontology.

### Antioxidant treatment corrects oxidative stress and restores neuron function

To assess whether oxidative stress was upstream or downstream of mitochondrial dysfunction, we treated AT32 mutant neurons with the antioxidant N-acetyl cysteine (NAC) for 10 days before oxidative stress levels were measured. NAC treatment reduced cytoplasmic CM-H_2_Xros signal by approximately 10% (Fig 7A, B) though treatment did not offer improvement in mitochondrial membrane potential (Fig 7C). AT32 corrected, mutant, and NAC treated mutant neurons were immunolabelled for tubulin to measure neurite extension, branching and survival (Fig 7D). NAC treatment significantly rescued neurite outgrowth in ATM deficient neurons, restoring the percentage FOV coverage of mutant neurons to levels comparable to corrected neurons Fig 7E). To assess if NAC and/or C7 treatment had any correctional effect on expression levels of the top genes dysregulated in AT neurons, RNA was isolated for gene expression analysis. *CHCHD2* mRNA was reduced in AT neurons treated with NAC compared to the untreated counterparts while *TUBB3* mRNA was unaltered (Fig 7F). *SOD3*, *NPAS4*, *SYP* and *SLC17A7* (VGLUT1) RNA expression in the treated AT neurons trended towards the corrected levels however did not reach levels of significance, indicating longer treatment periods may be necessary to correct these features (Sup Fig 10B).

**Figure 7:**
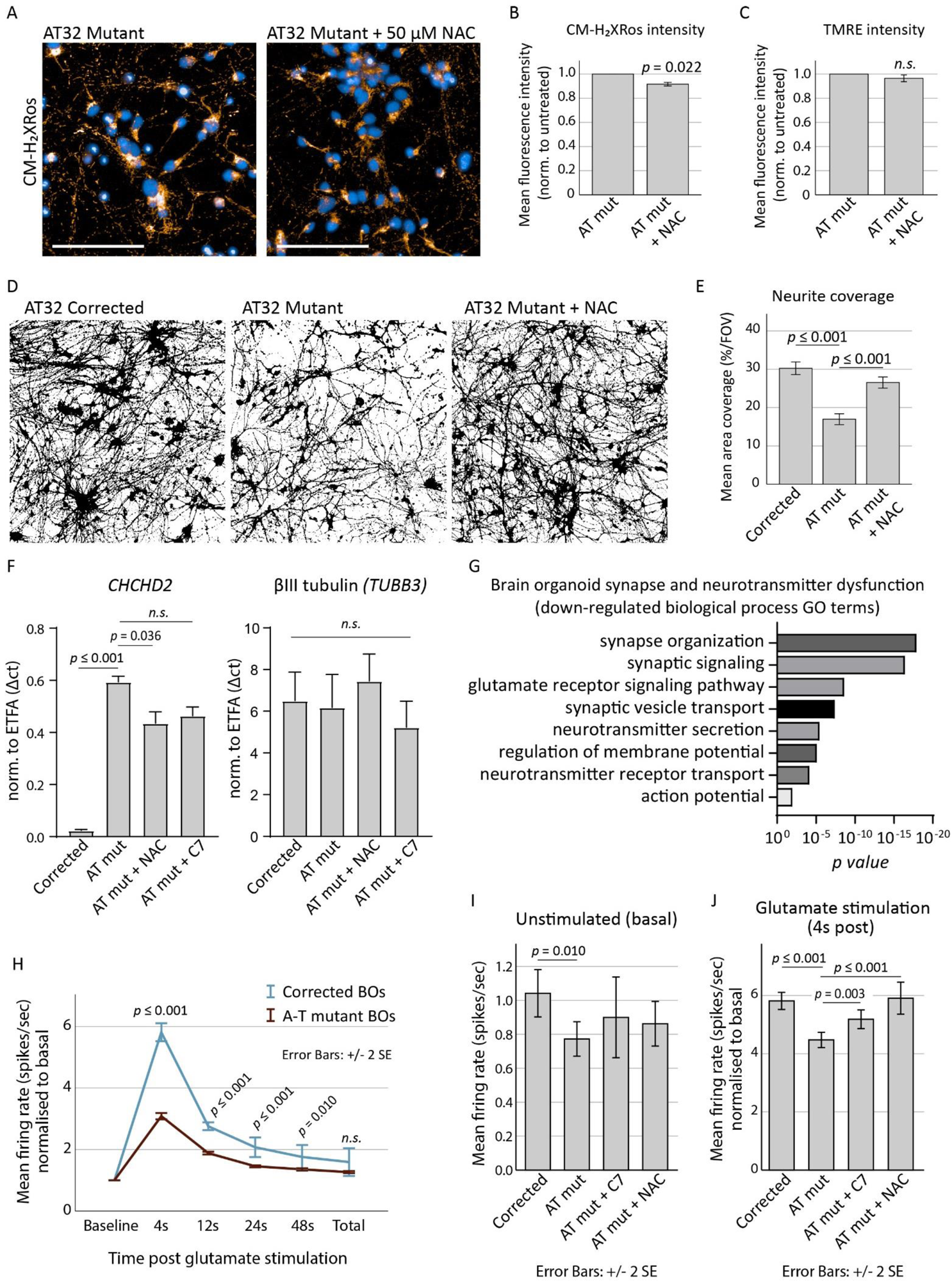
Antioxidant treatment reduces oxidative stress and restores neuronal firing. **A)** AT32 mutant neurons were treated with the antioxidant NAC for 10 days before oxidative stress levels were measured by CM-H_2_XRos. Scale bar 100 µm. **B)** CM-H_2_Xros fluorescence intensity was reduced following NAC treatment; t[3] = 4.388, *p* = 0.022, though TMRE fluorescence intensity **(C)** remained unchanged in the treated neurons; t(3) = 1.963, *p* = 0.072. Error bars mean ± 1SE. **D)** Representative binary βIII-tubulin immunochemistry of 3 independent AT32 corrected, mutant, and NAC treated mutant neuronal cultures. **E)** Binary βIII-tubulin immunochemistry was quantified to calculate the percentage coverage per FOV; F(2, 80) = 17.599, *p* ≤ 0.001, with Tukey post hoc analysis identifying a significant reduction (*p* ≤ 0.001) in mutant neurons compared to corrected, and a significant correction (*p* ≤ 0.001) of this phenotype with NAC treatment. Error bars mean ± 1SE. **F)** Total RNA was harvested from AT32 mutant and corrected neurons and used to quantify the mRNA expression levels of *CHCHD2* and *TUBB3.* Mutant neuronal cultures were treated with either NAC or C7. *ETFA* was used as housekeeper gene. ANOVA [F(5, 15) = 57.26, *p* ≤ 0.001] and Tukey post hoc analysis determined significant differences in *CHCHD2* mRNA levels, *p* values as indicated. *TUBB3* mRNA expression remained unchanged [F(5, 18) = 0.2079, *p* = 0.955]. 3 independent replicates, error bars mean ± SD. **G)** GO analysis conducted on up and down DEGs separately identified significant downregulation of pathways involved in synapse function and neurotransmitter signalling. **H)** Brain organoids were adhered to electrode chips and baseline electrical activity was recorded, followed by addition of glutamate, using an MEA platform. Mean firing rate (spikes per second) were calculated in time windows post stimulation with glutamate (4, 12, 24, 48 and total seconds post glutamate addition) and normalised to the basal firing rate. Student’s t test was used to determine significance, *p* values as indicated. Corrected organoids; 2834 active electrodes from 6 organoids on 4 MEA chips. Mutant; 4510 active electrodes from 5 organoids on 3 MEA chips. Error bars mean ± 2SE. **I-J)** AT32 mutant brain organoids were treated with C7 and NAC for two weeks before being adhered to electrode chips. Baseline electrical activity was recorded, followed by addition of glutamate. Mean firing rate (spikes per second) were calculated for baseline and for glutamate stimulation normalised to baseline. **(I)** Significance was determined by Welch’s ANOVA for baseline neuronal activity [F(3,5491) = 3.286, *p* = 0.020] and Games-Howell post hoc analysis found a significant reduction in baseline firing in mutant compared to corrected (*p* = 0.01). **(J)** Glutamate stimulation revealed significant differences between untreated and treated mutant organoids [F(3,6435) = 18.428, *p* ≤ 0.001, Games-Howell post hoc analysis as indicated]. Error bars mean ± 2SE. GO; gene ontology. MEA; multi-electrode array. NAC; N-acetyl cysteine. C7; heptanoate.

Given that our AT mutant neurons displayed reduced IEG expression, and KEGG and GO term analysis of organoid cultures showed altered synapse and neurotransmitter pathways (Figs 4 and 5, and highlighted in Fig 7G), we tested the neuronal firing capacity of 100-day-old organoids on a multi-electrode array platform, and determined if antioxidant treatments would correct any defects. We found that AT32 mutant organoids had a significant reduction in their response to glutamate stimulation (Fig 7H), and also showed a significantly reduced mean firing rate under resting (unstimulated) conditions compared to their gene corrected counterparts (Fig 7I). This reduction in the AT mutant organoids’ capacity to respond to glutamate aligns with the GO term transcriptome analysis that identified the glutamate receptor signalling pathway as significantly downregulated in the mutant organoids (Fig 7G). Brain organoids were treated with the antioxidant NAC as well as the anaplerotic agent heptanoate (C7), which replenishes TCA cycle activity and was shown to alleviate mitochondrial dysfunction, oxidative stress and cell death in ATM^-/-^ HBEC and AT patient-derived ONS cells under metabolic stress conditions [66]. Brain organoids treated with C7 and NAC did not show improvements in unstimulated neuronal firing rates (Fig 7I), however when exposed to glutamate stimulation, the NAC and C7 treated AT mutant organoids had mean firing rates that were significantly increased compared to the untreated AT organoids, and in the case of NAC treatment, the firing rate was restored to that of the gene corrected organoids (Fig 7J).

## Discussion

ATM is an important master regulator of mitochondrial homeostasis and oxidative stress as confirmed by studies in murine models [11–13] and in human patient cells [14–16]. Despite this, the extent of mitochondrial dysfunction and oxidative stress in patient neuronal cells remained uncharacterised. Here we, for the first time, utilise patient iPSCs to examine mitochondrial content, membrane potential and oxidative stress levels in human neural models across different stages of maturation, from stem cells and neuronal progenitors to 2D neuronal cultures, and finally, brain organoids. Our findings are of particular importance when considering AT neuropathology, especially given that current *in vivo* models do not manifest key neurological abnormalities, including neurodegeneration, that are consistently observed in individuals with AT [24].

We first interrogated the mitochondrial content, membrane potential and ROS production in the olfactory epithelial-derived ONS cells from 5 AT patients as compared to controls, observing no disease-specific differences. These findings are of particular interest, as previous studies showed that AT ONS cells possess neuronal-like features and exhibit hypersensitivity to radiation, defective radiation-induced signalling and cell cycle checkpoint defects [26], suggesting that impairments to these ATM-regulated DNA damage response pathways do not cause mitochondrial defects in this cell type. This aligns with the premise that mitochondrial impairments and oxidative stress in ATM deficient cells are not merely a downstream result of accumulating nuclear DNA damage [20, 67].

We next utilised iPSC derived from two of these AT patients to interrogate mitochondrial function over the course of neuronal maturation, from stem and progenitor cells through to brain organoids. We observed significant levels of mitochondrial impairments and oxidative stress that become progressively more pronounced with the increasing maturity of the neuronal model. Several ATM-dependent mechanisms that may contribute to neuronal degeneration in AT presented themselves, and these are discussed below. We further expanded our investigations to include aspects of aging which are tightly interwoven with mitochondrial health and function [62, 64], and documented significant levels of senescence in both cultured neurons and brain organoids. Lastly we employed an MEA platform as a functional readout of AT pathology, observing impaired neuronal function in brain organoids, which was alleviated by antioxidant treatment.

### AT patient neurons show deficits in mitochondrial membrane potential and increased oxidative stress

Mitochondrial membrane potential is generated by the proton pumps during oxidative phosphorylation and primarily reflects the activity of the electron transport chain, serving as an indicator of mitochondrial function and cellular energy status. We observed significant reductions in mitochondrial membrane potential in AT patient-derived neurons when compared to control/isogenic corrected neurons that became more pronounced over the course of the maturation process and is indicative of decreased mitochondrial function, respiratory chain activity and ATP synthesis. Supporting this is the reversal of CM-H_2_Xros accumulation within the mitochondria. While increased CM-H_2_Xros fluorescence in iPSC and NPC cultures were consistent with increased oxidative stress levels observed previously [32], their reversal to non-significance at 2 weeks and then decreased levels at 4 weeks was initially confounding. Mitochondrial sequestration of this dye is, however, dependent upon membrane potential and we concluded that the chronic loss of potential, combined with impaired mitochondrial homeostasis and mis-localisation, revealed in our neuronal models explained the decrease in CM-H_2_Xros intensity. Comparison of AT patient-derived neurons to control neurons treated with ATM inhibitors further supported this theory, as the acute ATM inhibition in neuronal models with otherwise healthy mitochondria did not reduce membrane potential and significantly increased CM-H_2_Xros accumulation in mitochondria, indicative of significant ROS production in the absence of a functional ATM protein, and consistent with previous studies demonstrating that these inhibitors also prevent ATM’s response to oxidative stress [20, 68]. Antioxidant treatment was shown to reduce oxidative stress in ATM deficient neurons and promote neurite growth and survival, however had no impact on membrane potential, consistent with oxidative stress being a downstream process following impaired mitochondrial function and/or membrane potential. Transcriptomic analysis of neuron and brain organoids further demonstrate a strong oxidative stress phenotype that is more pronounced in the organoids, suggesting that oxidative stress compounds with the maturity and complexity of the model.

### Processes of mitochondrial homeostasis, encompassing localisation, fission-fusion, mitophagy and biogenesis, are impaired in AT patient neurons and organoid models

Mitochondria are highly dynamic in nature, forming interconnected networks and actively migrating along neuronal processes to areas of high energy demand. They continuously engage in tightly controlled cycles of fission and fusion to maintain a functional population of mitochondria, underscoring the importance of mitochondrial health in the nervous system [69]. We observed disparity between AT neurons and control/isogenic corrected in terms of mitochondrial content and localisation within processes during the course of neuronal maturation, with reductions in soma mitochondrial presence coinciding with increases within neuronal projections at two weeks in culture, suggesting a shuttling of mitochondria to these processes, presumably as a compensatory mechanism to preserve neurite and synapse integrity. Neurons cultured up to 4 weeks no longer displayed this pattern, instead showing overall reductions. It is worth noting that while mitochondrial content diminished, the number of mitochondrial puncta within the neurites remained stable or increased, suggesting the lack of functional ATM compounds over time and leads to changes in the dynamics of mitochondrial architecture as they undergo fission, resulting in a fragmented mitochondrial network, as is observed in other neurodegenerative and age-related pathologies [61, 70] and correlating with observations by Luo, Lyu [37] who found ATM inhibition promotes excessive mitochondrial fission.

Recent findings identified compromised mitophagy and mitofission, as well as reduced NAD^+^/SIRT1 signalling downstream of PARP1 in ATM null mice and in ATM-deficient cells [54, 71]. NAD^+^ supplementation was found to improve mitophagy as well as DNA repair, and reduced the severity of AT neuropathology [54]. Interestingly, we observed in our AT patient neuronal models that both PARP1 and SIRT1 transcript and protein levels are reduced. PARP1 and SIRT1 are both nuclear enzymes and share NAD^+^ as a substrate, inhibiting each other’s activity; PARP1 activation causes rapid loss of mitochondrial potential and reactive oxygen species production while higher SIRT1 activity has roles in neuroprotection and metabolic homeostasis, improving mitochondrial function, promoting mitochondrial biogenesis, and protecting from ROS [60, 72, 73]. ATM and PARP1 are known to form a molecular complex *in vivo* in the absence of cellular damage [74]. In addition to direct interactions, ATM may also modulate the PARP/NAD^+^/SIRT signalling axis via modulation of NRF1, one of the key transcription factors for mitochondrial biogenesis genes. NRF1 is a phosphorylation target of ATM when activated by oxidative stress, but not DNA damage, and phosphorylation results in NRF1 nuclear translocation and upregulation of mitochondrial and OXPHOS genes. Human brain samples show NFR1 expression is enriched in Purkinje neurons compared to other cerebellar neurons, and nuclear translocation of NRF1 is reduced in AT patient brain samples [55].

Taken together, we hypothesise that ATM may regulate PARP/SIRT homeostasis by directly binding to PARP and/or via NRF1 modulation of PARP activity. NRF1 and SIRT1 activate *FOXO1* and *PGC-1α* to regulate mitochondrial homeostasis and biogenesis, transcript levels of which were all found to be disrupted in our AT patient brain organoids. In conjunction with disrupted PARP/SIRT signalling, dysregulation in some of the key signalling pathways responsible for fission (*OMA1, DRP1, SLP2*) and mitophagy (*PCG-1α, SQSTM1, PINK1* and *PRKN*) lead us to conclude that ATM is a master regulator of mitochondrial turnover that can impose its influence via multiple avenues, and that lack of functional ATM protein directly leads to impaired fission-fusion balance, increased biogenesis as well as reduced mitophagy, resulting in accumulation of damaged mitochondria.

Brain organoids generated from AT patients further demonstrate impaired mitochondrial maintenance, with AT organoids containing significantly more mitochondrial content than isogenic organoids, indicating that as well as impairments in mitochondrial transport and fission fusion, there are severe deficiencies with mitochondrial turnover and/or biogenesis. This is supported by our transcriptomic analysis, where we observed altered gene expression related to mitochondrial dynamics. Of particular note is the large upregulation in *CHCHD2* (*MNRR1*), a regulator of OXPHOS and of mitochondrial homeostasis [40]. Under stress conditions, mitochondrial import of CHCHD2 is inhibited, allowing accumulation in the nucleus where it functions as a transcription factor for a subunit of cytochrome c oxidase (*COX*) [39]. Importantly, overexpression of CHCHD2 was found to induce the mitochondrial unfolded protein response, autophagy, and mitochondrial biogenesis [41], aligning with the significant increase in mitochondrial content observed in our AT brain organoids. CHCHD2 can also act as an inhibitor of apoptosis [43, 44]; interestingly, many anti-apoptotic factors are upregulated during the process of senescence [75], which has presented itself as a distinguishing feature in AT brain organoids, as discussed below.

### AT patient neurons and brain organoids exhibit strong aging and senescence phenotypes

AT is now also considered a premature aging disease, with mitochondrial dysfunction as a key hallmark of the aging phenotype. Our lab has previously demonstrated that AT brain organoids display a strong senescence phenotype compared to hESC-derived control organoids, which could be rescued by inhibition of the cGAS-STING pathway [53]. Here we confirm this finding in 100-day-old isogenic AT and gene corrected brain organoids, as well as in 10-week-old 2D neuronal cultures, which showed increased SA-β-Gal, p16 and p21 staining. Mitochondrial dysfunction is often an under-unappreciated hallmark of aging and cellular senescence [62], and numerous studies indicate that impaired mitochondrial dynamics, such as decreased cycles of fission, contributes to the development of the senescence-associated secretory phenotype (SASP) and resistance to cell-death, with mitochondria becoming enlarged and hyperfused. Importantly, defects in mitophagy may also contribute to the induction of cellular senescence [64]. Aligning with our findings that PARP SIRT signalling and processes of mitophagy are disrupted, as well as observations of increased mitochondrial content, others have also found that NAD^+^ supplementation was able to promote mitophagy in AT fibroblasts, and the enhanced mitophagy prevented the development STING-mediated senescence in a PINK dependent manner [76]. NAD^+^ supplementation also inhibited neurodegeneration and senescence phenotypes in *ATM^−/−^* mice, and improved motor function and mitochondrial homeostasis. In light of these findings, and in combination with our observations of increased senescence in AT neuronal models, we conclude that mitochondrial dysfunction contributes to the aging phenotype of AT patients.

### Impaired neuronal signalling in ATM organoids is rescued by antioxidant or anaplerotic treatment

A lesser studied aspect of ATM is its role in intracellular vesicle and/or protein transport mechanisms, and its association with synaptic vesicles [77]. ATM binds to VAMP2 and SYNAPSIN-I, while ATM deficiency causes impaired cycling of synaptic vesicles, indicating a regulatory role for cytoplasmic ATM in neuronal activity [10]. ATM has since been more specifically located in excitatory (VGLUT1) vesicles [78]. Accordingly, reducing ATM in hippocampal neuronal cultures resulted in an excitatory/inhibitory imbalance toward inhibition, an increased number of GABAergic synapses, and reduced neuronal excitability [79]. Our findings strongly support these data, with GO and KEGG pathway analyses both indicating a reduction in synapse and vesicle transport terms, though it is probable that the significant mitochondrial impairments in our brain organoid models also contributes to downstream alterations in neuronal function. MEA analysis of 100 day old brain organoids demonstrates reduced levels of unstimulated/basal firing activity, and a reduced capacity of the neurons to respond to glutamate stimulation, which could be alleviated by NAC or C7 treatment. This is the first example of a functional neuronal defect in AT patient derived brain organoids, and indicates that while ATM may have mechanistic roles in synapse function which are unlikely to be countered by antioxidant treatment, these treatments may support neuronal activity by boosting mitochondrial health and reducing oxidative stress.

In conclusion, our data suggest AT patient neurons display progressive oxidative stress phenotypes, impaired mitochondrial membrane potential, dysregulated mitophagy, and impaired regulation of fission-fusion processes. We uncovered alterations in gene expression patterns related to maintenance of mitochondrial dynamics, including significant upregulation of CHCHD2 and altered PARP/SIRT signalling, which contribute to mitochondrial defects. Further, we identified increased levels of senescence and reduced neuronal activity in AT brain organoids. Our study demonstrates the complexity of mitochondrial dysfunction, and suggests that impairment of multiple ATM-dependent pathways involved in mitochondrial homeostasis and antioxidant signalling contribute to the neurodegenerative aspect of AT.

## Methods

Unless otherwise stated all reagents were purchased from Thermo Fisher Scientific, and cell culture mediums from Gibco (Thermo Fisher Scientific).

### Human olfactory epithelial cell culture

AT patients were diagnosed at the AT Clinic, University of Queensland Centre for Clinical Research, Brisbane, Australia. Biopsies from the olfactory mucosa of five AT patients and six healthy age matched controls were collected with informed consent as described previously [26] and the resulting olfactory epithelial (ONS) cells were cultured in DMEM/F12 supplemented with 10% foetal bovine serum (FBS), 1:100 GlutaMAX, 1:100 non-essential amino acids (NEAAs) and 1% Penicillin-Streptomycin (PenStrep). Cultures were passaged approximately every 5 days as they reached 70-80% confluency.

### Induced pluripotent stem cell culture and neuronal differentiation

Four iPSC lines were used in this study; AT and control ONS-derived iPSCs [29, 30], referred to as ‘AT’ and ‘control’, and AT32 patient and isogenic (gene corrected) iPSC pair [31–33]. The ATM mutations are described in Sup Fig 3. iPSCs were maintained in mTeSR Plus (STEMCELL Technologies) under feeder-free conditions on hESC-qualified Matrigel (Corning) coated plates. Clump passaging was performed using EDTA at 70-80% confluence approximately every 5 days. Stocks were cryopreserved as cell clumps using Synth-a-Freeze, and thawed in the presence of ROCK inhibitor Y-27632 (10 µM; Tocris).

iPSC lines were differentiated to neurons using a protocol modified from Shi, Kirwan [34]. Briefly, iPSC were cultured to 80% confluence as described above, then the medium rinsed with PBS and replaced with 3N neural differentiation medium (1:1 DMEM/F12 and Neurobasal medium, supplemented with 1:200 N2, 1:100 B-27, 1:100 GlutaMAX, 1:200 NEAAs, 2.5 µg/mL insulin, 50 µM β-mercaptoethanol and 1% PenStrep) supplemented fresh with 10 µM SB431542 (Milteny Biotec) and 0.1 µM LDN-193189 (Sigma). 3N neural differentiation medium with SB and LDN was replaced daily for 10 days, then cells were passaged using Dispase II (2.4 units/mL in HBSS) and re-plated at a 1:3 dilution on Matrigel coated plates in 3N neural differentiation medium. The neuro-epithelial cells were fed every other day, with 20 ng/mL FGF2 (PeproTech) added upon appearance of neural rosettes approximately 4 days following passage. Rosettes containing neural progenitor cells (NPCs) were manually selected and all subsequent passages were performed at 80% confluence using Accutase. NPCs were cultured either as 2D monolayers plated on Matrigel coated plates, or as 3D neurospheres in low attachment plates. For neuronal maturation, NPCs (P2 onwards) were seeded at 50,000 cells/cm^2^ on PLO and laminin coated plates or coverslips. 3N neural differentiation medium was supplemented with ascorbic acid (200 nM; Sigma), dibutyryl-cAMP (500 µM; Sigma), BDNF (20 ng/mL; PeproTech), GDNF (20 ng/mL; PeproTech) and DAPT (10 µM; Sigma). Neurons were cultured for between 2 and 10 weeks, and 50% culture medium was exchanged 3 times per week. For some assays, neuronal cultures were treated with ATM inhibitors KU-55933 or KU-60019 (Sigma, 10 µM each, 3 times per week with feeds), the antioxidant N-acetyl cysteine (NAC; 50 µM treated daily) or the anaplerotic agent fatty acid heptanoate (C7; 350 µM, 3 times per week with feeds). Treatments were commenced approx. 14 days post NPC plating on laminin and continued for 10-12 days prior to assessment at 4 weeks.

### Brain organoid generation

Brain organoids were generated from human AT32 mutant and gene corrected (isogenic pair) iPSCs. iPSCs were dissociated using Accutase and seeded at a density of 15,000 cells per well in 96-well low-attachment U-bottom plates (Sigma) in mTeSR Plus with 10 µM ROCK inhibitor. The 96-well plates were then spun at 330 g for 5 min to aggregate the cells. Spheroids formed overnight and were fed daily for 5 days with KSR medium (DMEM/F12, 20% Knock-out Serum Replacement (KSR), 1:100 GlutaMAX, 1:200 MEM-NEAA, Dorsomorphin (2 µM; StemMACS) and A-83-01 (2 µM; Lonza)). On day 6, the medium was changed to induction medium containing DMEM/F12, 1:100 N2, 1:100 GlutaMAX, 1:200 MEM-NEAA, 1 µg/ml heparin (Sigma), 1 µM CHIR 99021 (Lonza) and 1 µM SB-431542 (Sigma). On day 11 spheroids were embedded in Matrigel (15 µl droplets, gelled at 37℃ for 20 min) and transferred to low-attachment 24-well plates containing induction medium. From day 16, the medium was changed to organoid medium containing a 1:1 mixture of DMEM/F12 and Neurobasal medium supplemented with 1:100 N2 supplement, 1:50 B27 supplement, 1:100 GlutaMAX, 1:200 MEM-NEAA, 1% PenStrep, 50 µM 2-mercaptoethanol and 2.5 µg/mL insulin, with medium changed three times per week. Organoids were maintained in organoid media until the end of experiments, at 100 days post embedding.

### Immunochemistry

Cells were cultured in 96 well imaging plates (Costar; ONS, iPSC, NPCs) or PLO and laminin coated glass coverslips (for neurons). Cells were washed once with PBS prior to a 10 min fixation with 4% paraformaldehyde, permeabilised with Triton-X100 at 0.1% and blocked with 3% BSA for 1 hour. Primary antibodies were incubated overnight at 4℃. Secondary antibodies were incubated for 1 hour at room temperature. All antibodies used are listed in Supplementary Materials Table S1. Nuclei were counterstained with either Hoechst 33342 or DAPI prior to mounting and imaging. Neurite coverage experiments were conducted on coverslips and imaged using an Olympus BX61 microscope equipped with 20X air and 40X oil immersion. Captured images were analysed using FIJI (ImageJ). Channel intensity per field of view was measured using raw integrated density. Thresholds were used to create a binary image of the tubulin networks, then the percentage of pixels equal to 255 was measured using the area fraction function. Channel intensities and area fractions were normalised to the number of nuclei within each field of view.

### Mitochondrial function assays and Operetta high-content imaging

Total mitochondrial content was measured using MitoTracker™ Deep Red FM live cell stain (200 nM, incubated for 30-40 minutes). Mitochondrial membrane potential was measured using tetramethylrhodamine ethyl ester (TMRE; Abcam, 100 nM, 20 minutes), and ROS production was evaluated with MitoTracker™ Red CM-H_2_Xros, a chemically reactive reduced rosamine dye that only fluoresces once it enters an actively respiring cell and becomes oxidized by reactive oxygen species, and is then then sequestered in the mitochondria (400 nM, 30-40 minutes). Following incubation, neurons were rinsed with medium, counter stained with Hoechst 33342, rinsed again and returned to medium without phenol red for imaging. An Operetta CLS high-content analysis system (Perkin-Elmer) was used to acquire images. Generally, between 25 and 80 fields of view were acquired per well. At least 3 biological replicates were performed per experiment. All statistical analysis and interpretations were performed on well results, with unbiased analysis and quantification were performed using Harmony software (Perkin-Elmer).

### Unbiased image quantification using Harmony

Harmony software was used for unbiased automatic quantification of immunofluorescence and mitochondrial stains captured via the Operetta CLS high-content analysis system. Advanced flatfield correction was applied to all images. Analysis pipelines were modified from predefined Harmony algorithms and consisted of an input image, followed by determination of the nuclear region using Hoechst or DAPI staining (Find Nuclei), with modifications as required to diameter, splitting sensitivity and common thresholds. The cell body surrounding nuclear objects was identified (Find Cell Region) with modifications as required to thresholds. Cytoplasm regions were calculated by subtracting the nuclear region from the cell region. Cytoplasm and nuclear shape, area, and fluorescence intensity were used to exclude irregular or dead cells. Border objects were removed and fluorescence intensity per pixel unit of selected objects was calculated. For neuronal cultures, thresholds were set to restrict the cytoplasmic ROI to the perinuclear space. Spot detection algorithms were used to identify mitochondrial puncta along neuronal processes (Find Spots). For ONS, iPSC and NPCs, generally 25-40 fields of view per 96 well plate well were captured, with approx. 200-500 cells quantified per field of view. For neuronal cultures, between 50 and 80 fields of view were generally acquired per well of the 24 well plate wells, capturing between 200,000 and 400,000 cells, with the Find Spots algorithm generally identifying around 300,000 to 600,000 puncta per field of view. Field of view results were averaged and data output is provided as a whole well result (mean ± SD for each well).

### Senescence-associated β-galactosidase assay (SA-β-Gal)

Neuronal 2D cultures were washed in PBS, fixed for 10 minutes in 4% paraformaldehyde, washed, and incubated at 37°C (in the absence of carbon dioxide) with fresh SA-β-Gal stain solution (pH 6.0): Potassium ferricyanide 5 mM, Potassium ferrocyanide 5 mM, Sodium dihydrogen phosphate 0.4 M, Sodium hydrogen phosphate 92 mM, Sodium chloride 150 mM, Magnesium dichloride 2mM and 5-bromo-4-chloro-3-indolyl-β-D-galactopyranoside 1 mg ml^−1^. Staining was evident in 2-4 hours and maximal in 12-16 hours.

### Western blotting

Cells were rinsed with PBS and lysed with RIPA buffer (50 mM Tris pH 8, 150 mM sodium chloride, 1% Triton X-100, 0.5% sodium deoxycholate, 0.1% SDS), containing 1X complete protease inhibitor and 1X PhosSTOP phosphatase inhibitor (Roche). Genomic DNA in the lysate was sheared using 27G syringe needles. Protein concentration was estimated using a standard BCA assay. Samples were prepared at 30 µg of protein with DTT (100 mM) and 1X Laemmli SDS loading dye and heated at 65°C for 10 mins for samples for probing with total ATM. Samples for probing with rodent OXPHOS antibody cocktail were not heated due to MTCO1 (of complex IV) being very sensitive to heating. Lysates were resolved using denaturing TGS (Tris/glycine/SDS) buffer-based polyacrylamide gel electrophoresis (SDS-PAGE) followed by wet transfer (Tris/glycine/methanol) to nitrocellulose membranes. For the rodent OXPHOS antibody cocktail, transfer was instead to a PVDF membrane using high pH CAPS transfer buffer (10 mM CAPS [3-(Cyclohexylamino)-1-propanesulfonic acid], 10% methanol, pH 11). Membranes were rinsed in TBS-T (1X TBS, 0.05% Tween-20) and blocked in 5% skim milk powder in TBS-T for 1 hour. Primary antibody cocktails were diluted in blocking buffer and incubated at 4°C overnight. Membranes were washed and probed with HRP-conjugated secondary antibody for one hour at room temperature. Cross-reactivity was detected using Clarity ECL (BioRad) or Femto maximum sensitivity substrate (Thermo Fisher) for OXPHOS cocktail. Captured images were analyzed using Image Lab 4.1 (Bio-Rad, USA) software, and background adjusted values for bands of interest were used to calculate values for protein quantification. Protein expression levels were normalised to a loading control (either β-actin or α-tubulin) for each lysate.

### qPCR

Total RNA from cultured cells and brain organoids was isolated with RNeasy Mini Kit (Qiagen) according to the manufacturer’s instructions. 1 μg of total RNA was reverse transcribed using iScript cDNA Synthesis Kit (Bio-Rad). qPCR used PowerUp SYBR Green Master Mix (Applied Biosystems) on a Bio-Rad CFX96 Touch Real-Time PCR detection system. Each reaction was performed in duplicate with 3 biological replicates. Electron transfer flavoprotein subunit alpha (*ETFA*) or *GAPDH* were used as housekeeper genes for normalization. Primers sequences are listed in Table S2.

### RNA sequencing

AT32 mutant and gene corrected 2-week-old neurons (3 biological replicates each) and 100-day-old brain organoids (4 organoids each) were submitted for RNA sequencing. The integrity of RNA was confirmed by analysis on a 2100 Bioanalyzer RNA 6000 Pico Chip kit (Agilent) using the RNA Integrity Number (RIN) prior to sequencing by Novogene Ltd (Hong Kong). Total RNA libraries were generated using NEBNext® UltraTM RNA Library Prep Kit for Illumina® (NEB, USA) and were sequenced on an Illumina NovaSeq 150bp paired-end platform. Fastp was used to check quality on the raw sequences before analysis to confirm data integrity. Paired-end clean reads were mapped to the human genome assembly hg38 using HISAT2 software. Feature counts was used to count the read numbers mapped of each gene, including known and novel genes, and RPKM of each gene was then calculated based on the length of the gene and reads count mapped to this gene. Differential expression analysis between groups was performed using DESeq2 R package. The resulting P values were adjusted using the Benjamini and Hochberg’s approach for controlling the False Discovery Rate (FDR). Genes with an adjusted P (*padj*) value < 0.05 found by DESeq2 were assigned as a differentially expressed gene (DEG). Gene Ontology (GO term) enrichment and KEGG pathway analysis were conducted using the Database for Annotation, Visualization and Integrated Discovery [80] based on either up or down regulated DEGs. Brain organoid RNA sequencing data have been deposited in the European Nucleotide Archive with the primary accession code PRJEB72015.

### Multi-electrode array (BioCamX)

Multi-electrode array (MEA) analysis was conducted on organoids matured for at least 100 days using a high-resolution (4096 electrodes) BioCamX MEA platform (3Brain, Switzerland). Organoids were transitioned to BrainPhys medium with NeuroCult SM1 neuronal supplement (StemCell Technolgies) and were adhered to the BioCamX chip over 3 days prior to recording at 18kHz to capture electrical signals from single-cell spikes to population network events. Following acquisition, data was filtered using the integrated BrainWave5 software, and processed to identify spikes (defined as electrical activity >8 SD from the baseline) and bursts (5+ spikes with max intervals of 100ms). Spike frequency (mean firing rate; MFR), amplitude, duration and inter-event-intervals were calculated within BrainWave5. Electrodes not in contact with the organoid or recording a MFR of <0.05 spikes/sec were excluded from analysis. Generally, 2 organoids could be adhered to a chip and analysed separately using regions of interest to group channels. At least 3 separate recordings (with different organoids) were conducted. Recordings were initiated on stable chips (with saturated electrodes < 2-3%) and were captured over a 3 minute window, with glutamate (200 µM) applied at the 1 minute time point. MFR post glutamate treatments were normalised directly to the pre-glutamate (basal) MFRs. N-acetyl cysteine (NAC; 1 mM) or heptanoate (C7; 750 µM) treatments were applied for two weeks prior to recordings. BioCamX data is expressed as +/- 2 SE.

### Statistical Analysis

Unless otherwise stated, for each experiment at least 3 biological repeats (N, referring to differentiations conducted from the point of iPSCs) were conducted with a minimum of 2 technical repeats (*n,* referring to parallel wells or differentiations conducted simultaneously from the same NPC culture). Unless otherwise indicated, data were presented in bar graphs as the mean ± 1 standard error (SE) or using box and whisker plots, representing the median and interquartile (IQ) range, with whiskers representing the minimum and maximum data points. Statistical significance was determined by t-test (independent samples) or one-way ANOVA and post-hoc analysis was carried out using Tukey HSD. In cases where Levene’s test for homogeneity showed a significant effect of variance, the more stringent Welch’s One-Way ANOVA was carried out and Games-Howell post-hoc test was used to reduce the chance of type 1 errors. The *p* values were reported as calculated to three decimal points. Analysis of data was conducted using IBM SPSS Statistics 27 or GraphPad Prism.

## Acknowledgements

We thank Maria Kasherman and Shaun Walters from the School of Biomedical Sciences Microscopy and Image Analysis Facility (The University of Queensland) for technical support, and all E.W. laboratory members for discussions. We extend our sincere gratitude to the patients and their families for donating tissue, and for their enthusiastic support for our research.

## Author contributions

J.A. performed staining and analysis shown in Figure 6E, F, and prepared brain organoids for RNA sequencing; C.G.I. generated the results in Figure 6G-I; H.C. generated the human brain organoids; A.F. assisted with sequencing analysis; Z.H. assisted with iPSC generation, differentiation and culture maintenance; and M.L. and A.M.S. provided the ONS cells and intellectual support. H.L. generated all other data, designed the study, and wrote the paper. E.W. assisted with study design and intellectual support. All authors edited the paper.

