## Supplementary data for "Ataxia Telangiectasia patient-derived neuronal and brain organoid models reveal mitochondrial dysfunction and oxidative stress"

**Table S1. Antibodies used in this study.**

| Antibody/antigen | Application details | Host | Catalogue No | Manufacturer |
| --- | --- | --- | --- | --- |
| **PRIMARY ANTIBODIES** |  |  |  |  |
| **ATM - 2C1(1A1)** | 1:1000 (WB)<br>*Corresponds to C-terminus of ATM protein (2577-3056 aa)* | Mouse | ab78 | Abcam |
| **NESTIN** | 1:500 (IF) | Mouse | 33475 | CST |
| **PAX6** | 1:250-1:500 (IF) | Mouse | SC-81649 | Santa Cruz |
| **βIII tubulin** | 1:1000 (IF) | Mouse | T8578 | Sigma |
| **MAP2** | 1:500 (IF) | Mouse | MAB3418 | Merck |
| **MAP2** | 1:500 (IF) | Rabbit | PA5-17646 | Thermo Fisher Scientific |
| **NANOG** | 1:2000 (IF) | Mouse | 4893 | CST |
| **NANOG** | 1/400 (IF) | Rabbit | 4903 | CST |
| **SOX2** | 1:400 (IF) | Rabbit | 23064 | CST |
| **OCT4** | 1:100 (IF) | Mouse | MAB4419 | Millipore |
| **TRA-1-81** | 1:200 (IF) | Mouse | MAB4381 | Millipore |
| **TRA-1-60** | 1:200 (IF) | Mouse | MAB4360 | Millipore |
| **DCX** | 1:500 (IF) | Mouse | SC-271390 | Santa Cruz |
| **ZO1** | 1:500 (IF) | Mouse | 33-9100 | Thermo Fisher Scientific |
| **PARP** | 1:1000 (WB) | Rabbit | 9532 | CST |
| **SIRT1** | 1:1000 (WB) | Rabbit | 2496 | CST |
| **α-tubulin** | 1:5000 (WB) | Mouse | T5168 | Sigma |
| **MTCH2** | 1:500 (IF) | Rabbit | ab113707 | Abcam |
| **Total OXPHOS rodent antibody cocktail**<br>(reacts with human) | 1:1000 (WB)<br>This OXPHOS cocktail contains 5 abs:<br>**CI: NDUFB8**, ab110242, 20kDa<br>**CII: SDHB**, ab14714, 30kD<br>**CIII: UQCRC2**, ab14745, 48kDa<br>**CIV: MTCO1**, ab14705, 40kDa, and,<br>**CV: ATP5A**, ab14748, 55kDa | Mouse | ab110413 | Abcam |
| **β-actin** | 1:10,000 (WB) | Mouse | MCA5775GA | Bio-Rad |
| **p21** | 1:500 | Mouse | 2946 | CST |
| **p16** | 1:500 | Rabbit | 80772 | CST |
| **CHCHD2** | 1:100 | Rabbit | NBP2-92335 | Novus Biologicals |
| ***SECONDARY ANTIBODIES*** |  |  |  |  |
| **Anti-rabbit IgG, HRP** | 1:3000 | Goat | 7074 | CST |
| **Anti-mouse IgG, HRP** | 1:3000 | Horse | 7076 | CST |
| **Anti rabbit IgG 488** | 1:500 | Goat | A11008 | Thermo Fisher Scientific |
| **Anti-rabbit IgG 568** | 1:500 | Donkey | A10042 | Thermo Fisher Scientific |
| **Anti-rabbit IgG 568** | 1:500 | Goat | A11011 | Thermo Fisher Scientific |
| **Anti-rabbit IgG 647** | 1:500 | Goat | A21245 | Thermo Fisher Scientific |
| **Anti mouse IgG 488** | 1:500 | Goat | A11001 | Thermo Fisher Scientific |
| **Anti mouse IgG 568** | 1:500 | Goat | A11004 | Thermo Fisher Scientific |
| **Anti mouse IgG cy3** | 1:500 | Donkey | 715-165-151 | Jackson ImmunoResearch |

IF; immunofluorescence. WB; western blot. CST; Cell Signalling Technology

**Table S2. qPCR primers used in this study.**

| **Target gene** | **Forward** | **Reverse** |
| --- | --- | --- |
| *CHCHD2* | CGAGGCCTGACATCACTTACC | TTTGCAAGTCGGCACTGT TTC |
| *TUBB3* | GCAACCAGATCGGGGCCA | GTT CATGATGCGGT CGGGAT |
| *SOD3* | CTCTCTTGGAGGAGCTGGAAAG | GTACATGTCTCGGATCCACTCC |
| *NPAS4* | GTGAGGCTACAGGCCAAGAC | AGGGCAGCATGGTCGGAGTG |
| *SYP* | CTCGGCTTTGTGAAGGTGCT | CTGAGGTC ACTCTCGGTCTTG |
| *SLC17A7*<br>(*VGLUT1*) | CTGGGGCTACATTGTCACTCA | GCAAAGCCGAAA ACTCTGTTG |
| *ETFA* | TGTTGATGCTGGCTTTGTTC | TGGATGGCTCCAGATATTCC |

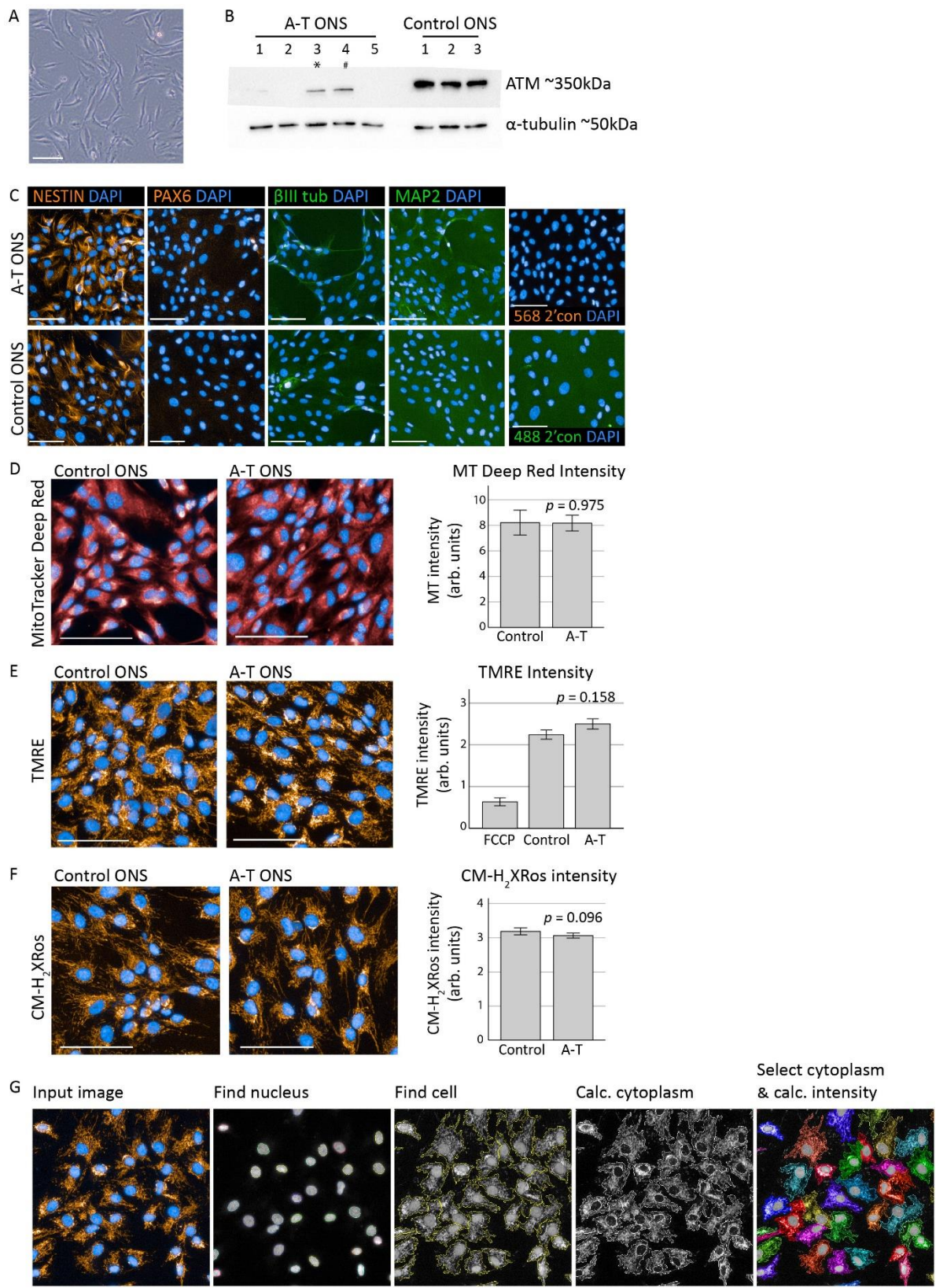

**Supplementary Figure 1: Characterisation and mitochondrial assessment of AT ONS cells**

ONS cells were cultured as monolayers (A) for all experiments. (B) Western blot analysis of ATM expression in whole cell extracts from 5 AT ONS lines compared to control (wild type). α-tubulin used as loading control. AT ONS line 3 (indicated by an asterisk) was reprogrammed to iPSCs, while AT ONS line 4 (hash symbol) is derived from the same patient (AT32) whose fibroblasts were previously reprogrammed. (C) ONS cells (passages 8-12) were assessed by immunochemistry for intermediate filament protein NESTIN (~80% positive), neural progenitor marker PAX6 (negative), and mature neural markers βIII tubulin and MAP2 (negative). (D-F)

Mitochondrial content, membrane potential, and oxidative stress levels were assessed in AT and control ONS cells by high content imaging and automatic quantification of mitochondrial stains. 5 control and 5 AT ONS cell lines were used. Total mitochondrial content was assessed by MitoTracker Deep Red (**D**; $t(7) = 0.033$, $p = 0.975$), mitochondrial membrane potential was evaluated by TMRE (**E**; $t(8) = 1.558$, $p = 0.158$, with FCCP used as a negative control) and reactive oxygen species (ROS) production was measured using MitoTracker Red CM-H<sub>2</sub>Xros (**F**; $t(18) = 1.759$, $p = 0.096$). Scale bars are 100 $\mu\text{m}$, error bars are mean ± 1 SE. **G**) Automated image analysis by Harmony software was used to quantify MitoTracker Deep Red, TMRE and CM-H<sub>2</sub>Xros fluorescence intensity captured on an Operetta high-content analysis system. The analysis pipeline was modified from a predefined Harmony algorithm and consisted of an input image, followed by ‘find nucleus’ and ‘find cell region’ steps. Cytoplasm region was subsequently calculated by subtracting the nucleus from cell regions. Nuclear and cytoplasmic regions were filtered using thresholds to remove dead cells and border objects, fluorescence intensity per pixel unit of selected objects was calculated and results were produced as averages per well. Object counts and cell areas were also obtained for quality control purposes. ONS; human olfactory epithelial cells. FCCP; carbonyl cyanide-p-trifluoromethoxyphenylhydrazone.

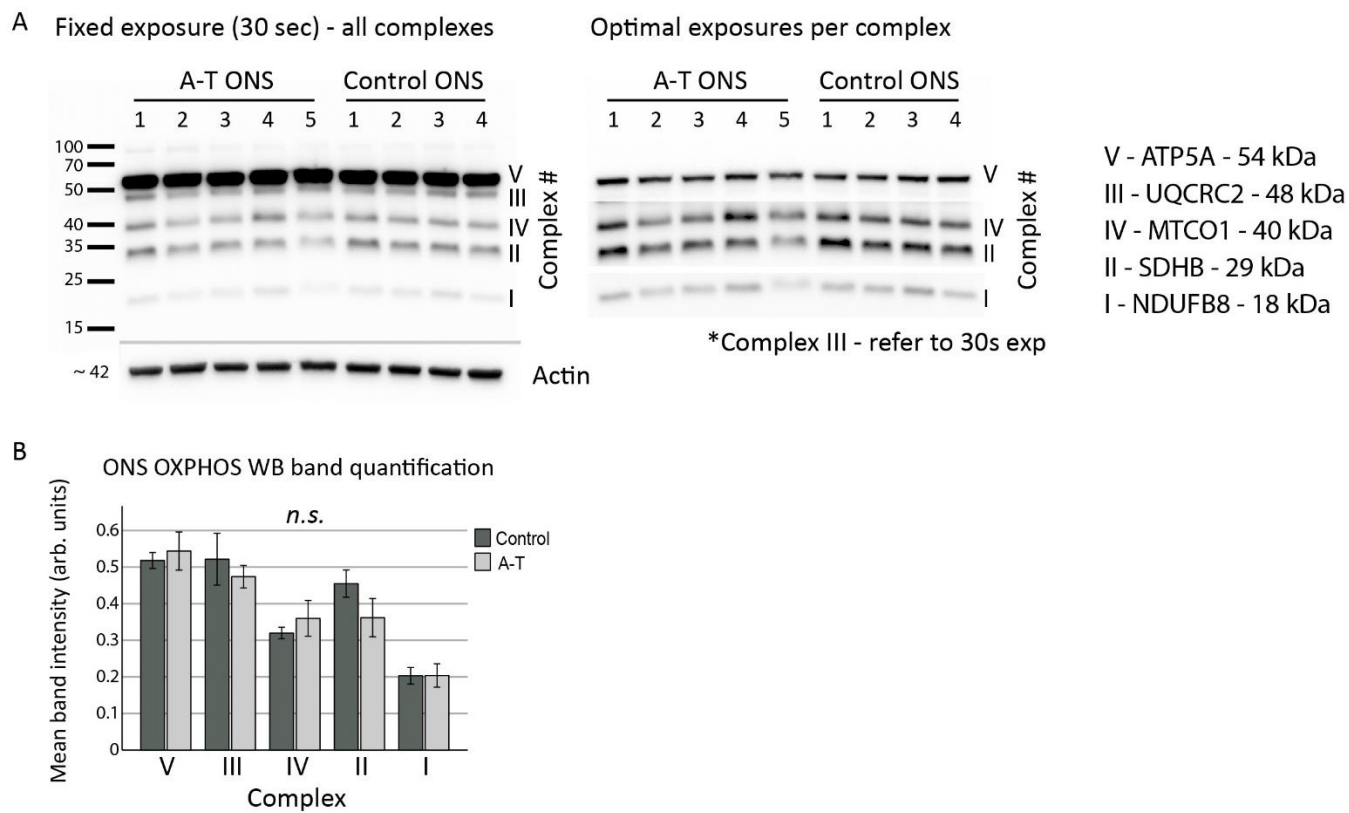

### Supplementary Figure 2: Assessment of OXPHOS complex expression levels in ONS cells

**A)** Relative levels of individual oxidative phosphorylation complexes were quantified by western blot using a total OXPHOS antibody cocktail containing antibodies against ATP5A (complex V, ~ 54 kDa), UQCRC2 (complex III, ~ 48 kDa), MTCO1 (complex IV, ~ 40 kDa), SDHB (complex II, ~ 29 kDa) and NDUF8 (complex I, ~ 18 kDa). A fixed exposure image was captured demonstrating presence of all five complex bands, followed by optimal exposures per complex band. Optimal exposure images were subsequently used to quantify expression levels after normalisation to actin loading control (**B**). Band intensity quantification showed no significant differences between control and AT ONS cells. 4 control and 5 AT ONS cell lines were used. Error bars are mean ± 1 SE.

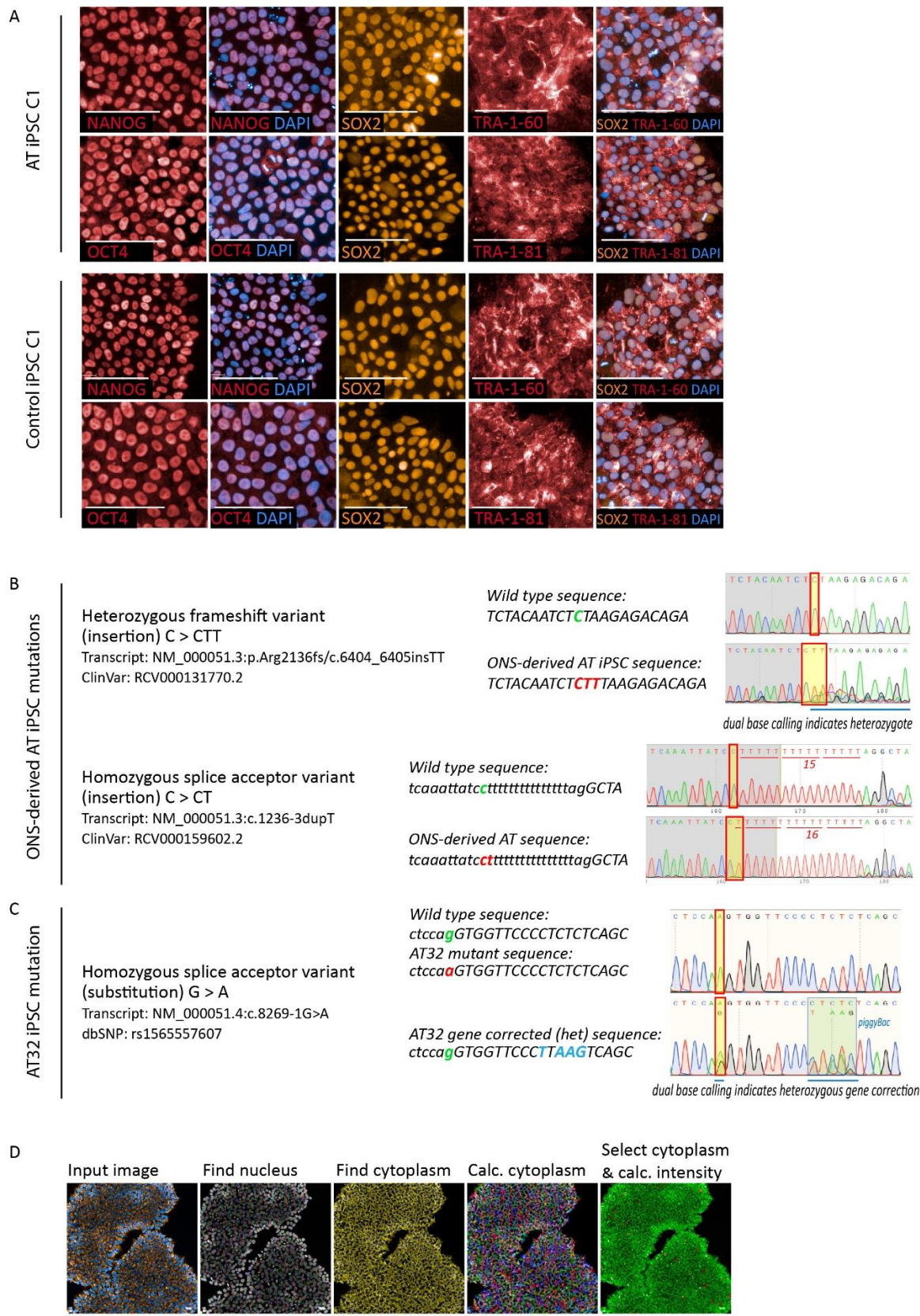

**Supplementary Figure 3: AT patient-derived iPSC lines retain ATM mutations and display equal pluripotency markers**

**A)** iPSCs from AT iPSCs (ONS-derived) robustly express NANOG, OCT4, SOX2, TRA-1-60 and TRA-1-81 at similar levels to control ONS-derived iPSCs. Scale bar indicates 100 μm. **B-C)** Mutations in the ATM gene were initially identified during patient diagnosis and were confirmed using Sanger sequencing following iPSC generation and at the commencement of this project. **B)** ONS-derived AT iPSCs contains compound mutations in exons 10 and 44. Exon 10 contains a homozygous splice acceptor variant (insertion/duplication; C>CT; NM\_000051.3:c.1236-3dupT; ClinVar RCV000159602.2), and exon 44 contains a heterozygous frameshift variant (insertion; C>CTT; NM\_000051.3:p.Arg2136fs/c.6404\_6405insTT; ClinVar RCV000131770.2). **C)** AT32 iPSCs contain a homozygous splice acceptor variant (substitution; G>A; NM\_000051.4:c.8269-1G>A) which has previously been gene corrected to a heterozygous status using a PiggyBac transposase. Mutations are highlighted in red, while the PiggyBac is identified in blue. **D)** Automated image analysis by Harmony software was used to quantify MitoTracker Deep Red, TMRE and CM-H<sub>2</sub>Xros fluorescence intensity in iPSCs. The analysis pipeline was modified from a predefined Harmony algorithm and consisted of a ‘find nucleus’ and ‘find cytoplasm’ regions. Nuclear and cytoplasmic regions were filtered using thresholds to remove dead cells and fluorescence intensity of selected objects was calculated. Results were output as averages of the whole well. Object counts and cell areas were also obtained for quality control purposes.

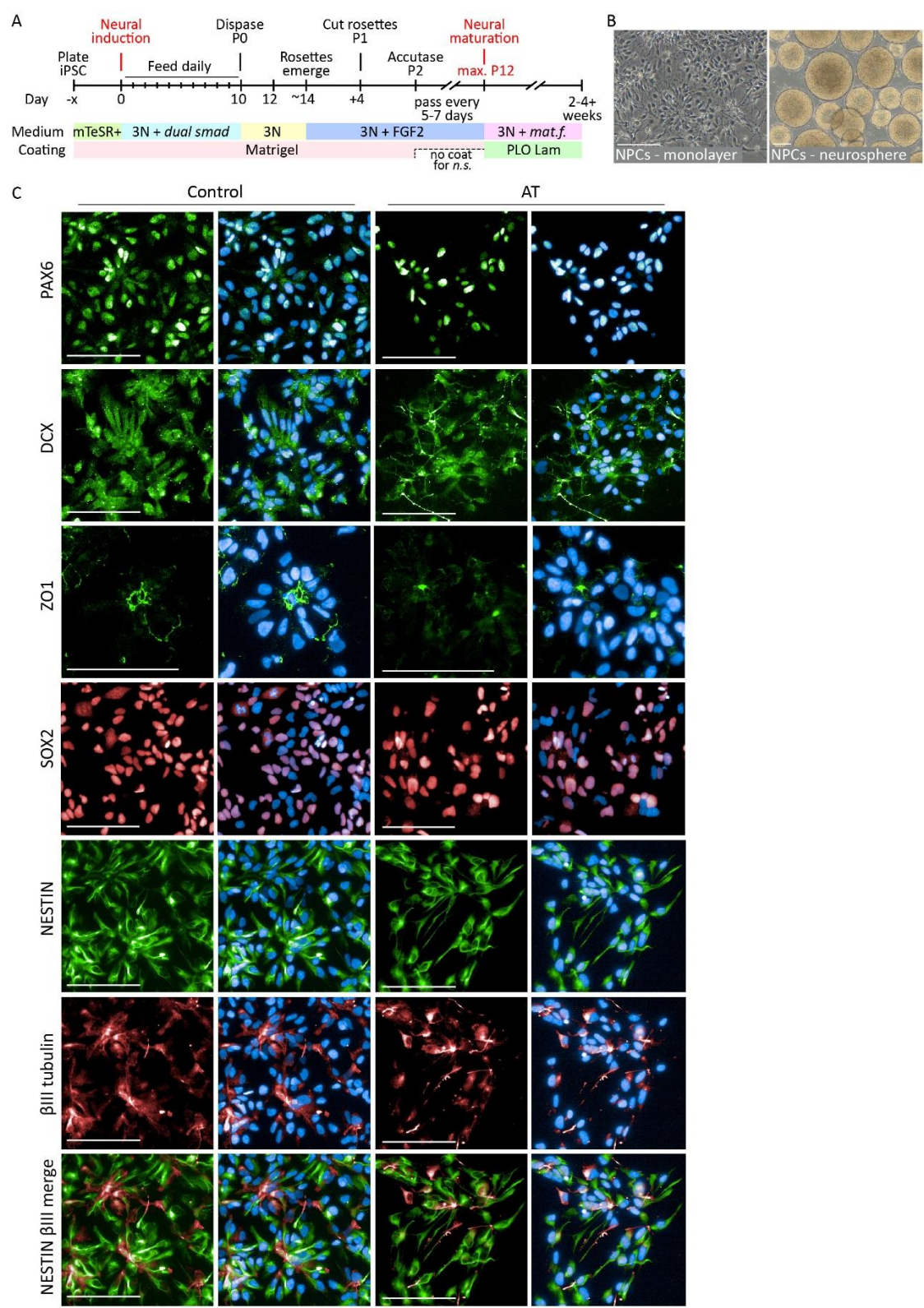

**Supplementary Figure 4: Differentiation and immunocytochemistry characterisation of iPSC-derived NPC cultures**

**A)** iPSCs were differentiated to NPC and neuron cultures in a protocol modified from Shi et. al. 2012. **B)** Example morphologies of NPC cultures as either 2D monolayers or as neurospheres. **C)** Control and AT NPCs were characterised to ensure no significant differences in differentiation capacity. Rosette-selected NPCs were positive for progenitor markers PAX6, SOX2, DCX and NESTIN. ZO-1 identified the basal membrane of NPCs in rosette formations, and βIII tubulin positive progenitors comprised approximately 20% of both AT and control cultures. Scale bars indicate 100 μm.

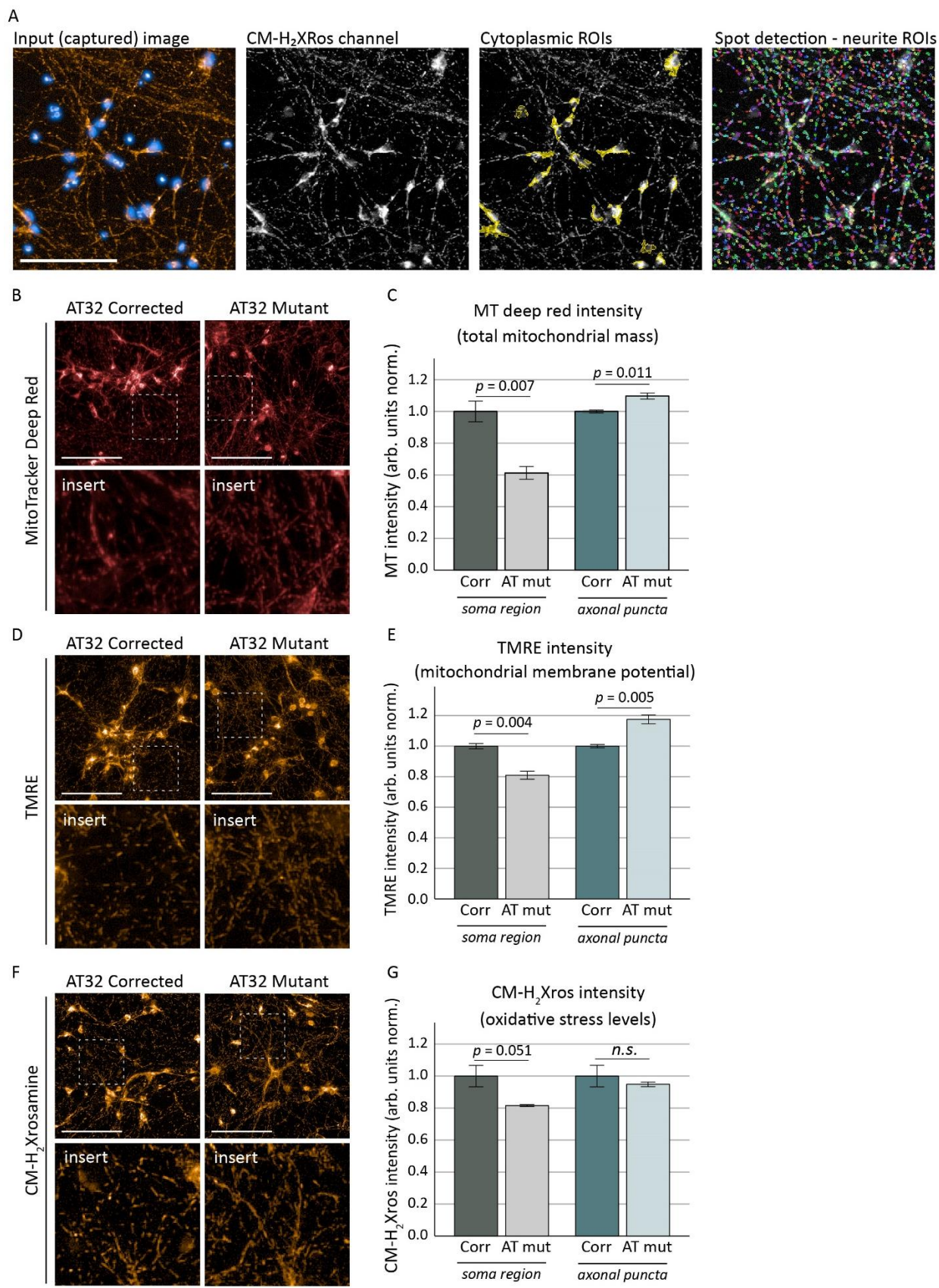

**Supplementary Figure 5: Image quantification and mitochondrial assessments on AT patient iPSC-derived 2-week-old neurons**

**A)** An Operetta was used to image MitoTracker Deep Red, TMRE and CM-H<sub>2</sub>Xros fluorescence intensity in 2- and 4-week neuronal cultures, and quantification was subsequently carried out in Harmony using an automated image analysis pipeline. The pipeline consisted of an input image, followed by ‘find nucleus’, ‘find cell region', and 'calculate cytoplasm' steps as described in Sup Fig 1. Thresholds were set to restrict the cytoplasmic ROI to the perinuclear space, henceforth describes as the soma region. Spot detection was used to identify mitochondrial puncta along neuronal processes, and intensity and puncta size thresholding was set to exclude perinuclear staining. Dead cells and border objects were removed and fluorescence intensity per pixel unit of selected objects was calculated. **(B-G)**. Mitochondrial content and membrane potential in AT32 mutant and gene corrected neurons at 2 weeks *in vitro* were assessed by MitoTracker Deep Red and TMRE, while oxidative stress was measured by CM-H<sub>2</sub>Xros in 3 biological replicates and quantified in both the soma region and in neuronal processes. MitoTracker Deep Red **(B)** identified significant differences in mitochondrial content and localisation between AT32 mutant and gene corrected neurons **(C)**. Soma intensity was reduced ($t(4) = 5.053$, $p = 0.007$) while neurite mitochondrial content was increased ($t(4) = -4.531$, $p = 0.011$). TMRE staining **(D, E)** reflected MitoTracker observations, with peri-nuclear mitochondria having a reduced membrane potential ($t(4) = 6.100$, $p = 0.004$) and peripheral mitochondria having an increased potential ($t(4) = -5.759$, $p = 0.005$). Oxidative stress levels as measured by CM-H<sub>2</sub>Xros **(F, G)** were not significantly different at 2 weeks *in vitro* ($t(4) = 2.763$, $p = 0.051$ and $t(4) = 0.749$, $p = 0.495$ for soma and neurite intensity respectively). Scale bars; 100 $\mu\text{m}$. Error bars are mean ± 1 SE.

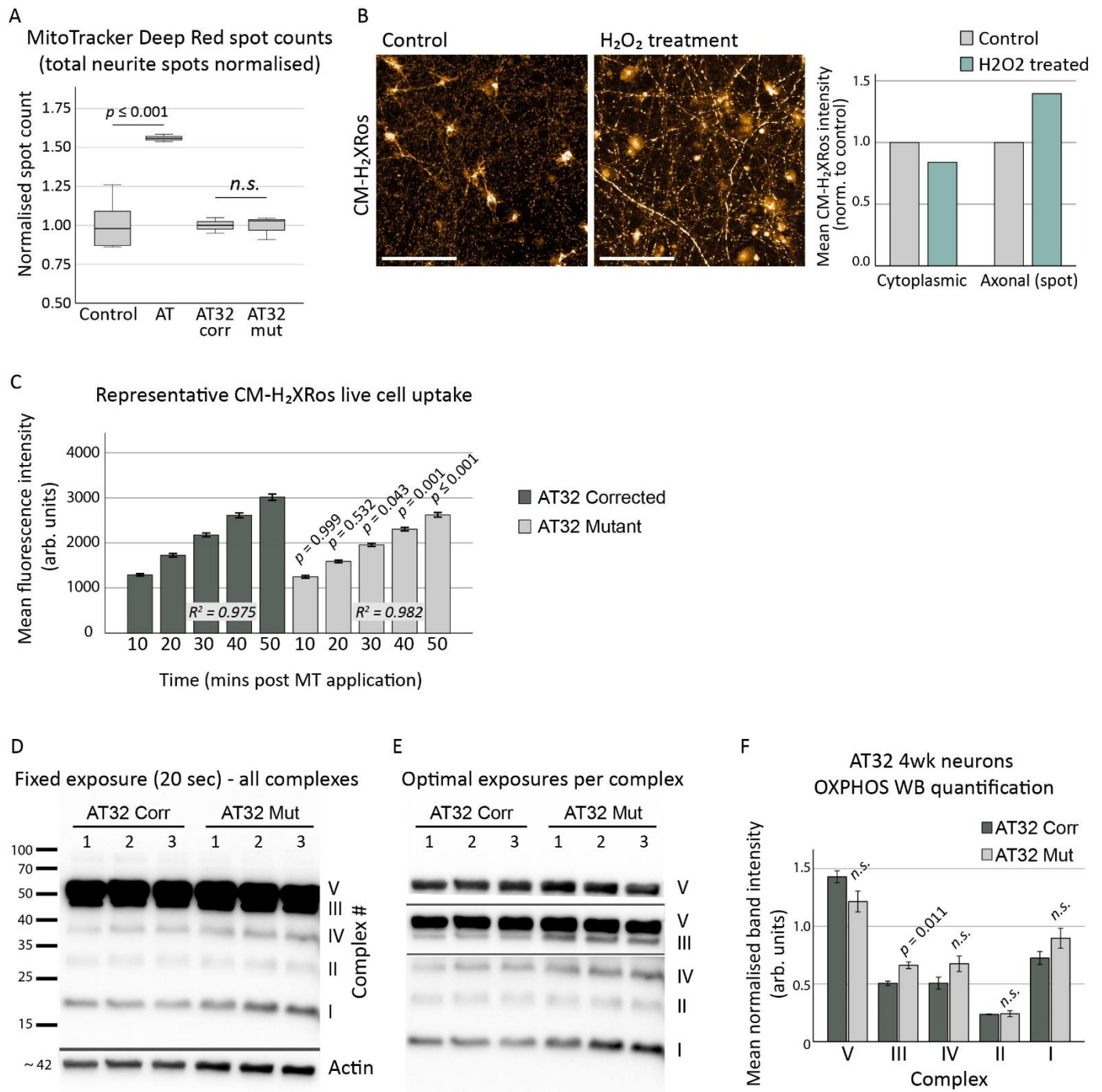

**Supplementary Figure 6: Additional analysis of mitochondrial stains and OXPHOS complex quantifications in AT patient iPSC-derived neurons**

**A)** Total spot count (neurite mitochondrial puncta detected by MitoTracker Deep Red staining) showed a significant increase in AT neurons [$t(9) = -6.546$, $p \leq 0.000$] compared to control, while AT32 isogenic neurons were consistent across genotype [$t(4) = 0.103$, $p = 0.923$]. **B)** Control neurons treated with H<sub>2</sub>O<sub>2</sub> as a positive control confirmed increased fluorescence of CM-H<sub>2</sub>Xros dye in mitochondria in the context of oxidative stress. Scale bar; 100 μm. **C)** Representative experiment of CM-H<sub>2</sub>Xros live cell dye uptake over time, confirming a linear sequestration of CM-H<sub>2</sub>Xros to mitochondria independent of genotype. An ANOVA test detected significant differences between the AT32 isogenic corrected and AT32 mutant lines [$F(9, 44) = 179.614$, $p \leq 0.000$], with Tukey HSD post hoc analysis $p$ values as indicated, and $R^2$ values of 0.975 and 0.982 respectively. **D, E)** Protein levels of individual OXPHOS complexes in 4-week-old neurons were detected by western blot using a total OXPHOS antibody cocktail. A single exposure image (**D**; fixed 20 sec) and optimal exposure images (**E**) for each complex band were captured. Band densities were measured and normalised to actin loading to quantify protein expression levels. Complex III protein levels were significantly increased in AT32 mutant neurons [$t(4) = -4.496$, $p = 0.011$], though other complexes failed to reach levels of significance (**F**). Error bars are mean ± 1 SE.

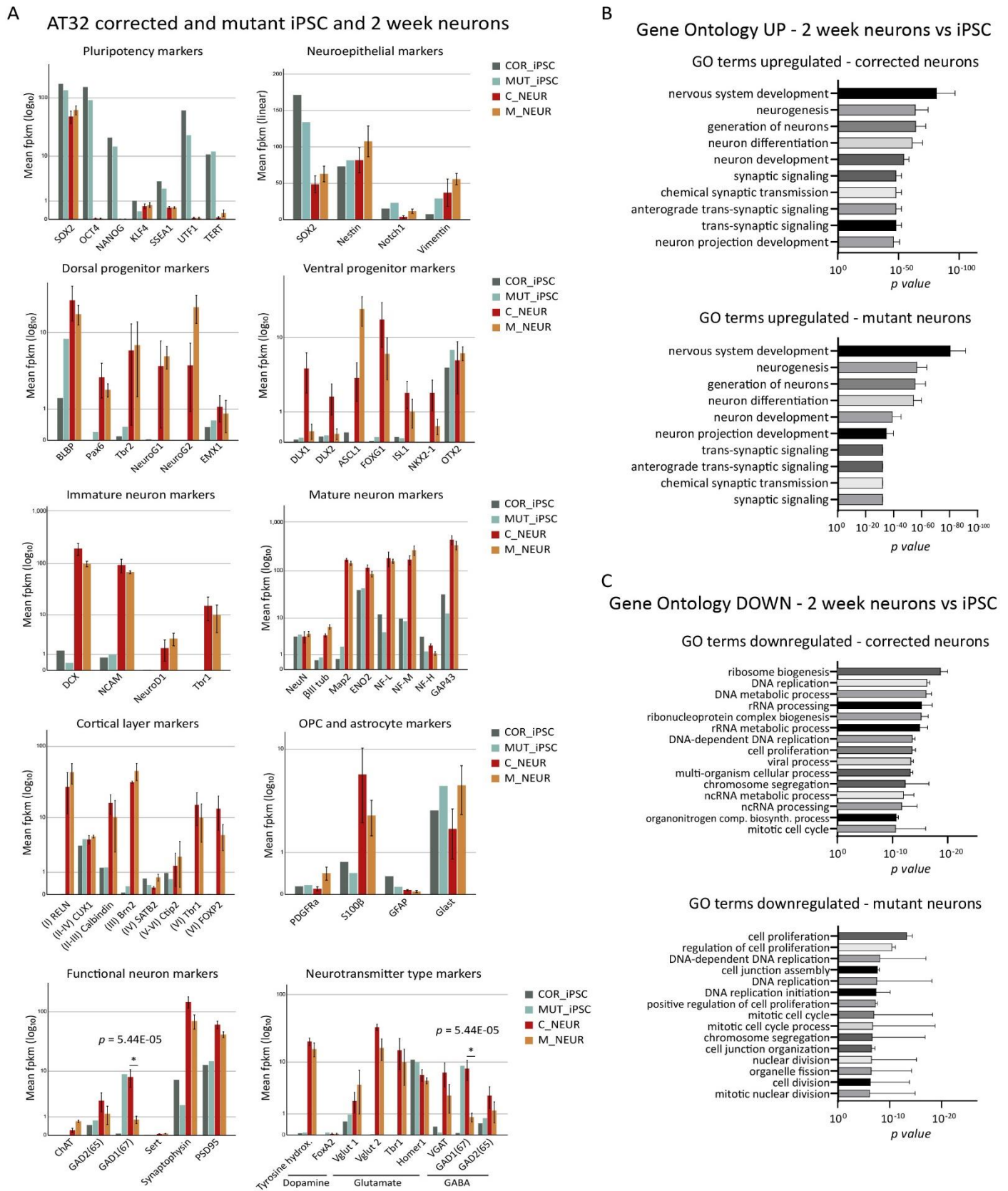

**Supplementary Figure 7: Analysis of differentiation potential in AT32 mutant and corrected 2-week-old neuronal cultures**

**A)** FPKMs were compared between iPSC and 2-week-old cortical neurons for AT32 mutant and gene corrected lines to ensure differentiations were consistent between genotype and replicates. Markers of pluripotency, neural progenitors (neuroepithelium, dorsal and ventral progenitor markers), and neuronal cells (immature and mature neuronal markers, cortical layer markers, oligodendrocyte and astrocytic markers, as well as markers of neuronal function and neurotransmitter type were all assessed. In agreement with our immunocytochemistry data, no significant differences were observed between control and AT for any of the investigated markers, except for *GAD1* (a GABAergic neuron marker), which was reduced in AT32 mutant neurons compared to the gene corrected lines ($p = 5.44\text{E-}05$). **B-C**) GO terms were calculated separately based on the upregulated and downregulated DEGs in neurons compared to the undifferentiated iPSCs. **B**) Upregulated GO terms were the same between ATM corrected and mutant neurons compared to their respective iPSCs while down regulated GO terms (**C**) were consistent with maturing neurons exiting the cell cycle and becoming post-mitotic. 3 biological replicates for AT32 mutant and corrected neurons and 1 replicate of iPSCs were sequenced. Error bars are mean ± 1 SE. COR/C; AT32 gene corrected; MUT/M; AT32 mutant. DEG; differentially expressed gene. GO; gene ontology.

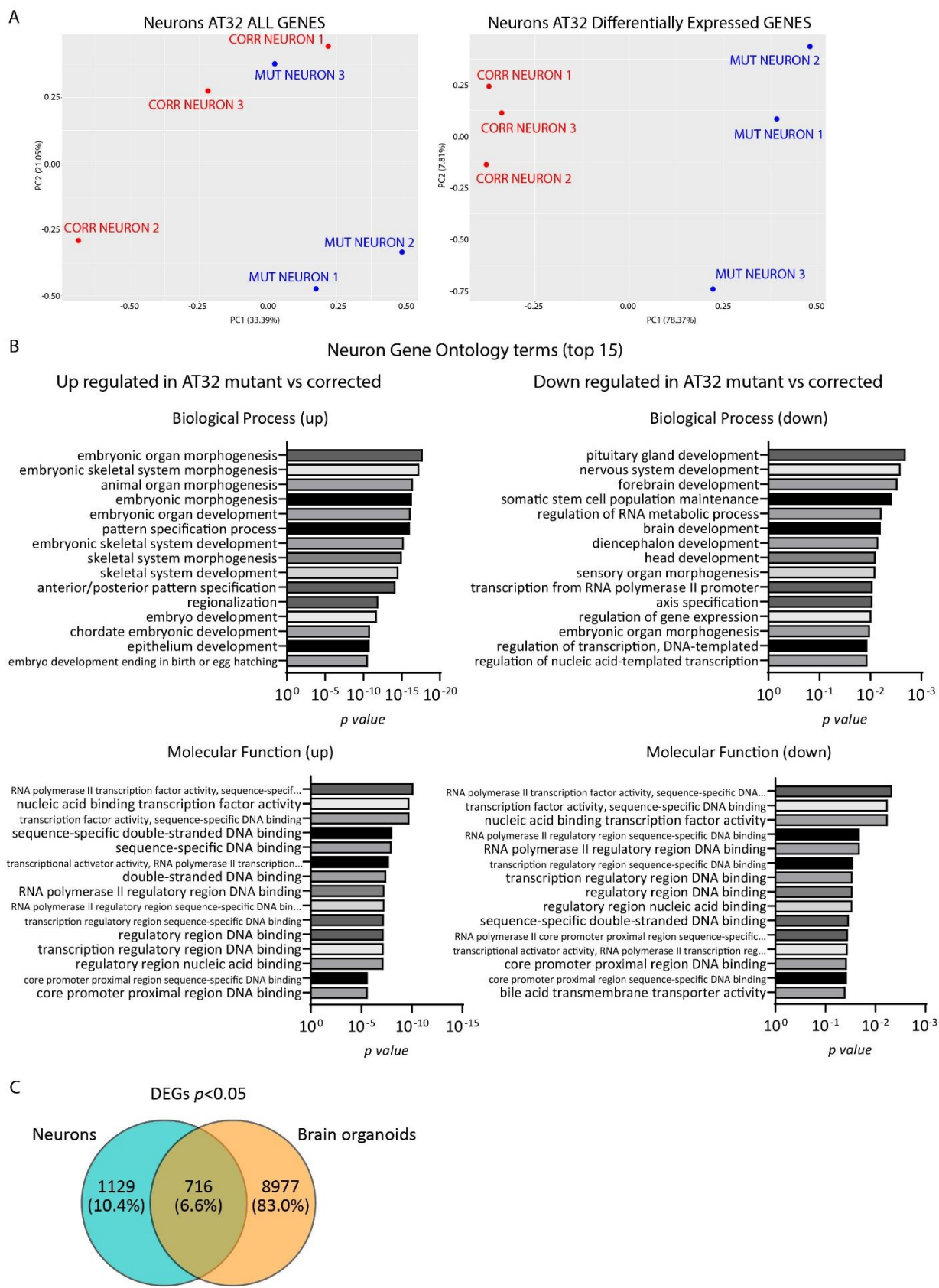

**Supplementary Figure 8: Additional transcriptomic analysis of AT32 2-week-old neuron cultures**

**A)** PCA were conducted using both all gene and DEG data sets, with disease specific separation of neurons identifiable by PC1 in the DEG data set. **B)** The top 15 GO terms for Biological Process and Molecular Function calculated using up-regulated and down-regulated DEGs separately. **C)** Venn diagram depicting overlap in genes between neuron and brain organoid data sets. PCA; principle component analysis. DEG; differentially expressed gene. GO; gene ontology.

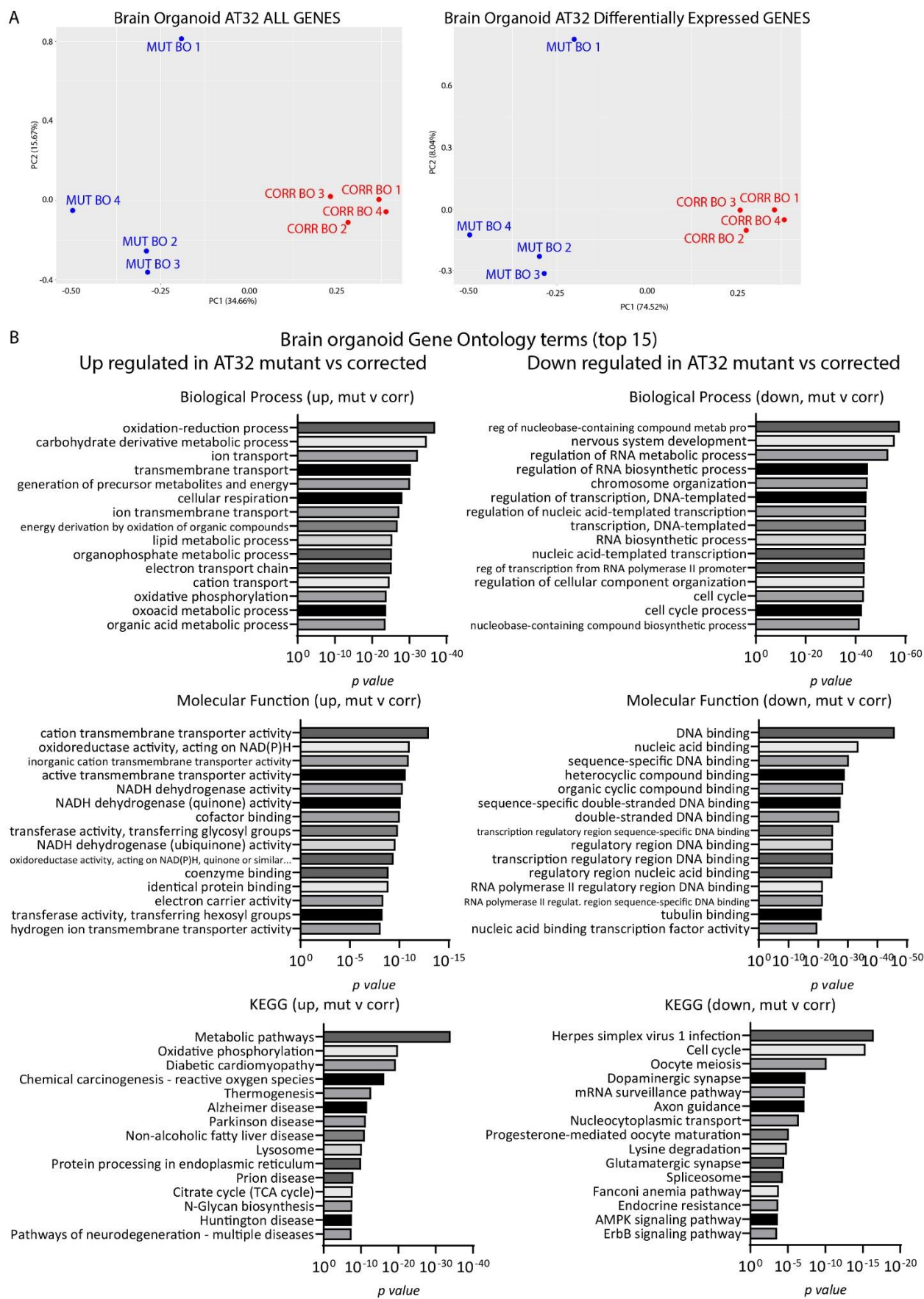

**Supplementary Figure 9: Additional transcriptomic analysis of AT32 brain organoid cultures**

**A)** PCA were conducted using both all gene and DEG data sets, with disease specific separation of brain organoids identifiable by PC1. **B)** The top 15 GO terms (Biological Process and Molecular Function) and KEGG pathways were calculated using up-regulated and down-regulated DEGs separately. BO; brain organoid. PCA; principle component analysis. DEG; differentially expressed gene. GO; gene ontology.

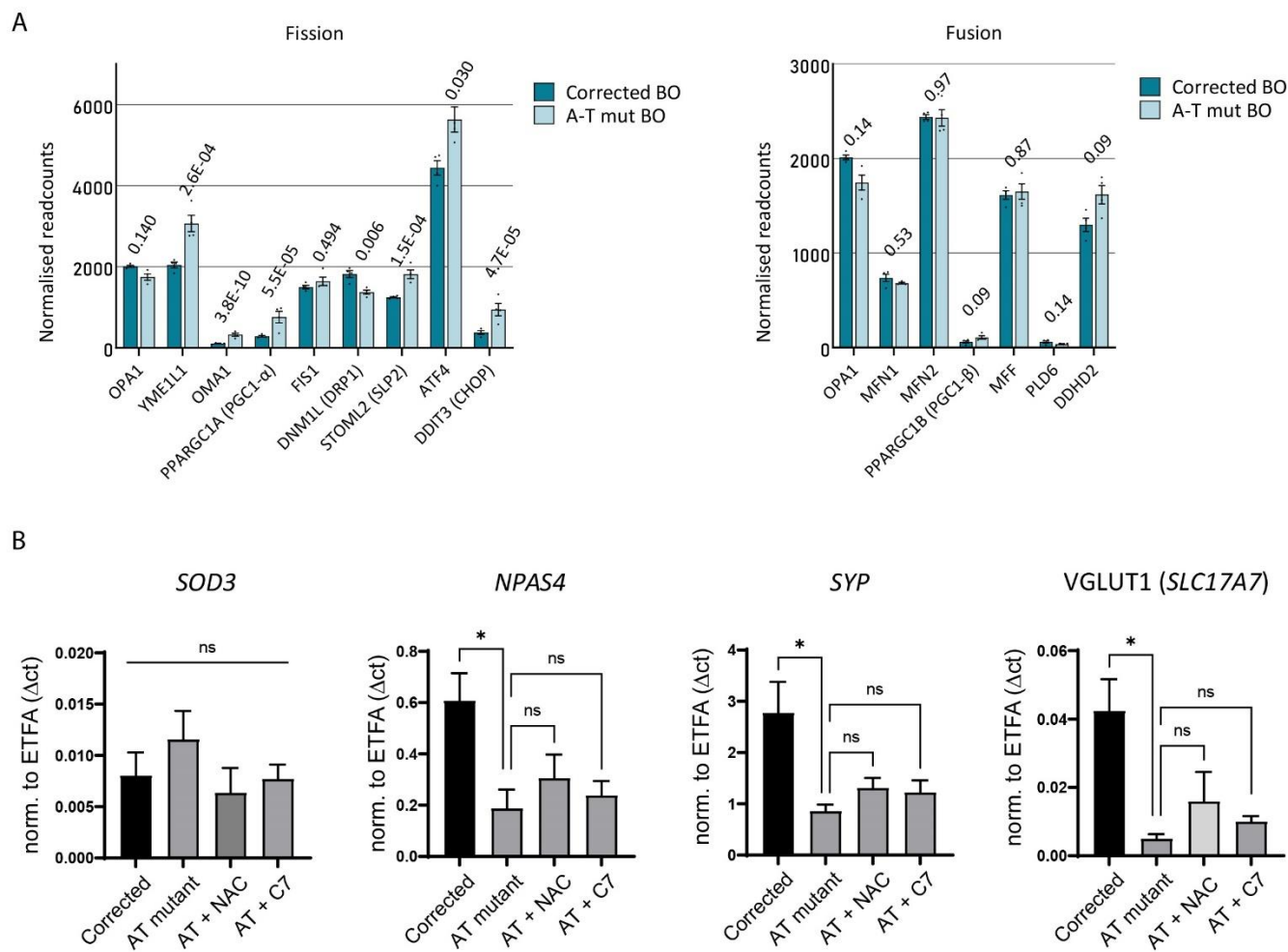

#### Supplementary Figure 10: Extended mRNA expression analysis

**A)** Normalised readcounts of an extended list of genes involved in fission and fusion process from AT32 brain organoid RNA sequencing data. *padj* values as indicated. **B)** mRNA expression in AT32 neurons treated with antioxidant (NAC) or anaplerotic (C7) agents. Total RNA was harvested from neuronal cultures and used to quantify the mRNA expression levels of *SOD3*, *NPAS4*, *SYP* and *SLC17A7* (VGLUT1) by qPCR. Mutant neuronal cultures were treated with either NAC or the anaplerotic agent heptanoate (C7). *ETFA* was used as normalizer gene. ANOVA determined no significant effect of treatment compared to untreated mutant neurons across 3 independent biological samples. Error bars mean ± SD. NAC; N-acetyl cysteine. C7; anaplerotic agent heptanoate.
